# Evolutionary constraint predicts the repeatability of transcriptional divergence during repeated adaptation

**DOI:** 10.64898/2026.09.28.755181

**Authors:** Zachary J.C. Tobias, Diana J. Rennison

## Abstract

The repeated evolution of similar phenotypes in response to shared ecological pressures provides striking examples of natural selection and raises broader questions about the predictability of evolution. Evolutionary constraint may shape this predictability, while the apparent extent of parallelism may also depend on the biological scale at which it is measured. When are adaptive solutions channeled along the same route? We addressed this question using three sympatric benthic-limnetic ecotype pairs of threespine stickleback (*Gasterosteus aculeatus*), characterizing gene expression in gill and liver from wild-caught fish. We tested whether proxies of pleiotropic and coding-sequence constraint predicted the repeatability of transcriptional divergence across lakes. At the gene level, differential expression was strongly biased toward parallel rather than antiparallel responses. At the functional level, numerous pathways showed parallel enrichment, but the conservation of their underlying member-gene responses varied substantially, indicating that similar functional outcomes can arise with differing degrees of gene-level concordance. Most importantly, several proxies of evolutionary constraint predicted transcriptional repeatability, particularly in gill: broadly expressed, highly connected, and more slowly evolving genes showed more stable expression responses across lakes, with similar relationships evident for repeated use of genes within functionally parallel pathways. Together, these results show that both gene-level constraint and biological scale contribute to variation in molecular repeatability and suggest that the predictability of repeated adaptation may reflect an interaction between genetic architecture, population-genetic context, and the level of biological organization at which parallelism emerges.

## Introduction

A central goal of evolutionary biology is to understand when evolution is predictable. Independent populations exposed to similar selective pressures often evolve similar phenotypes (Blount et al. 2018; Bolnick et al. 2018), yet the molecular changes underlying these outcomes range from strikingly repeatable to highly idiosyncratic (e.g. Cooper et al. 2003; Colosimo et al. 2005; Jones et al. 2012; Elmer et al. 2014; Fischer et al. 2021). Explaining this variation in molecular repeatability is critical for understanding whether evolutionary outcomes reflect a broad landscape of interchangeable solutions or are instead channeled along a limited number of viable trajectories.

Evolutionary constraint is likely to be an important determinant of this predictability. Genes differ substantially in their evolutionary degrees of freedom (Pál et al. 2006; Futuyma 2010; Lindblad-Toh et al. 2011), because changes to their sequence or regulation differ in their potential consequences elsewhere in the organism, for example through pleiotropy and/or epistasis (Yeaman et al. 2018; Mackay and Anholt 2024). Broadly expressed genes (Watanabe et al. 2019), highly connected components of regulatory networks (Promislow 2004; He and Zhang 2006), and genes under strong sequence/structural constraint (Franzosa and Xia 2009; Echave et al. 2016) may therefore have fewer viable routes of evolutionary change (Zhang 2023). Although such constraints are generally expected to limit evolutionary responses, restricting the set of viable changes could also make those responses more repeatable, and thus predictable.

The apparent predictability of molecular evolution, and its relationship to evolutionary constraint, may depend on the level of biological organization at which it is measured (Bolnick et al. 2018; Lau and Oakley 2021). Independent populations need not recruit the same mutations, genes, or regulatory changes to achieve similar phenotypic or functional outcomes. At the same time, distinct gene-level responses may converge on the same cellular processes or pathways, producing greater repeatability at higher levels of organization than is apparent from individual genes. Molecular parallelism is therefore not a single property of an adaptive response, but can emerge at multiple hierarchical levels, from individual genetic changes to gene-expression programs and broader biological functions. Distinguishing among these levels is necessary both to characterize how predictable molecular evolution actually is, and to identify the mechanisms that shape this predictability.

Gene expression is a particularly useful level at which to examine evolutionary predictability because it occupies an intermediate position between genetic variation and organismal phenotype. Similar changes in expression can arise from distinct underlying regulatory changes, allowing transcriptional parallelism to emerge even when evolution is not repeated at the nucleotide level. At the same time, expression responses can be compared both among individual genes and across the functional pathways to which they contribute, revealing whether similar higher-level outcomes arise through repeated recruitment of the same molecular components or through alternative transcriptional routes. Gene expression therefore provides a natural framework for asking not only how repeatable molecular evolution is, but at what level of biological organization that repeatability emerges. In natural populations, however, repeated expression differences measured in wild-caught individuals may integrate heritable regulatory divergence with environmentally induced responses associated with divergent ecologies. Thus, transcriptional parallelism can be viewed as repeatability of the expression phenotype without necessarily implying reuse of the same underlying genetic mechanisms. Many studies, in both the lab and field, have documented repeatable shifts in transcription across populations or species adapting to similar conditions and/or with repeated phenotypic change (Hart et al. 2018; Jacobs et al. 2020), while others have demonstrated idiosyncratic shifts (Fischer et al. 2021). Of course, in reality transcriptional repeatability is not a binary property, but rather a continuum of the extent to which the same genes respond in the same direction and magnitude across independent evolutionary transitions, with many individual studies documenting both parallel and nonparallel components of gene expression evolution (Manousaki et al. 2013; Hanson et al. 2017; Velotta et al. 2017). This variability in the magnitude of repeatability allows us to examine which properties of genes, and aspects of the regulatory architecture, determine the degree to which expression responses are repeated across independent evolutionary replicates.

One potentially key determinant of evolutionary constraint is pleiotropy, whereby a single genetic or regulatory change affects multiple phenotypes. Changes to gene expression differ in their potential pleiotropic consequences depending on factors such as whether a gene is tissue-specific versus broadly expressed or peripheral versus highly embedded in a regulatory network. The classic theoretical expectation is that genetic or regulatory changes with fewer pleiotropic consequences should be favored during adaptation, as a means of mitigating off-target deleterious consequences (Fisher 1930; Orr 2000). More recent empirical and theoretical work suggests that intermediate or even high levels of pleiotropy may be favored in the specific context of parallel evolution, particularly when selection acts on correlated traits in the same direction, such that a single pleiotropic change can move multiple traits toward their adaptive optima simultaneously (Rennison and Peichel 2022; Battlay et al. 2024). These investigations have mostly considered parallelism at the genomic level; there are comparatively few studies that have probed this relationship specifically at the level of gene expression (but see McGirr and Martin 2018; Durkin et al. 2024; Lai et al. 2025; Ramirez-Lanzas et al. and others).

The existing literature provides contrasting empirical examples. Durkin et al. (2024) found that genes with more tissue-specific expression, and thus ostensibly lower pleiotropy, were overrepresented among transcriptionally parallel genes during repeated colonization of colder northern environments by house mice. Lai et al. (2025) observed that genes with broader expression across tissues, and thus *greater* estimated pleiotropy, were more likely to be parallel in their expression response during experimental evolution of heat tolerance among replicate lines of *Drosophila simulans*. A third study on parallel evolution of trophic specialization in Bahamian pupfish found no relationship between expression parallelism and a gene’s degree of pleiotropy, as estimated by the number of protein-protein interactions, associated gene ontology terms, and developmental stages across which it is expressed (McGirr and Martin 2018). Together, these contrasting results suggest that the relationship between pleiotropic constraint and transcriptional repeatability is highly context dependent and remains unresolved.

We investigate evolutionary repeatability using the independently derived sympatric threespine stickleback (*Gasterosteus aculeatus*) ecotypes. In British Columbia, there are three lakes where an open water-specialist ecotype (limnetic) and a shore-specialist ecotype (benthic) have evolved in sympatry (Figure 1). These ecotypes differ in a suite of ecologically relevant traits, including morphology, physiology, and behavior (McKinnon and Rundle 2002). For decades these ‘ecotype’ or ‘species’ pairs have been the focus of intense study on speciation in the face of gene flow and the genetic basis of parallel evolution (Schluter and McPhail 1992; Conte et al. 2015; Schluter et al. 2025). Nonetheless, to date there has been limited consideration of how the ecotypes differ in their patterns of gene expression (but see Rennison et al 2016, Keagy et al 2024) or the extent of parallelism among lakes.

**Figure 1.**
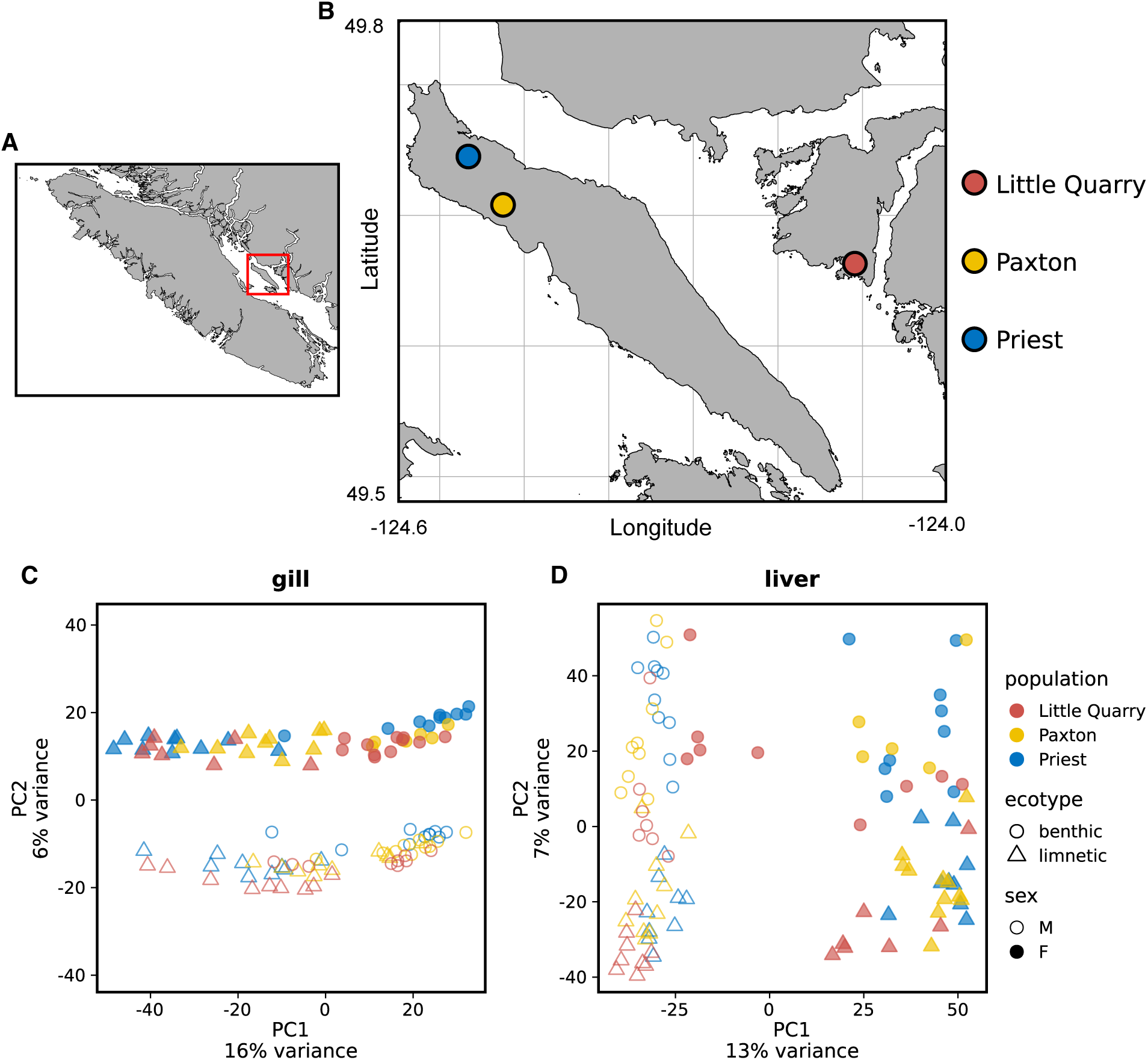
A) Extent of study area (in red square) in relation to Vancouver Island and mainland British Columbia B) Map showing Texada Island (with Paxton Lake and Priest Lake) and Nelson Island (with Little Quarry Lake). C-D) PCAs of transcriptome-wide gene expression data. Each point represents an individual fish’s expression profile. Color indicates lake, shape indicates ecotype, and fill indicates sex. In gill, samples cluster strongly by ecotype along PC1 and sex along PC2. In liver, samples cluster strongly by sex along PC1 and ecotype along PC2. Five benthic females from Little Quarry cluster with benthic males across lakes.

We characterized whole transcriptome expression for the three replicate benthic-limnetic pairs in two tissues, gill and liver, to identify the primary functional axes of ecotypic divergence. Although expression divergence was strongly biased toward parallel rather than antiparallel responses across lakes, the degree of repeatability varied substantially among genes and pathways. This variation allowed us to address an unresolved question raised by mixed prior results: do features associated with pleiotropic and coding-sequence constraint systematically predict which transcriptional responses are repeatedly recruited during parallel adaptation? We tested this question using multiple proxies of evolutionary constraint, asking whether they were associated with both transcriptome-wide stability and gene reuse within parallel functional pathways, and whether these relationships reflected the repeatability rather than simply the magnitude of expression divergence.

## Methods

### Sampling, RNA extraction, and sequencing

Fish were collected from Little Quarry Lake, Paxton Lake, and Priest Lake in April and May, 2024 using minnow traps under British Columbia Fish Collection Permit NA24-845596. Site and sample information is given in Table 1. After collection, fish were held with aeration at ambient temperature before dissection. Individuals were euthanized with MS-222 (500 mg/L) and the liver and right gills were dissected and placed in RNAlater. Tissue samples were stored overnight at 4 °C to allow the RNAlater to perfuse the tissue, followed by -20 °C for long-term storage.

**Table 1.** Summary of sampling and BRBseq reads per individual across lakes, ecotypes, and tissues.

| **Tissue** | **Lake** | **Ecotype** | **Female n** | **Male n** | **Total n** | **Mapped reads, mean**<br>**+/- SD (M)** |
| --- | --- | --- | --- | --- | --- | --- |
| Gill | Priest | Benthic | 10 | 10 | 20 | 3.88 +/- 0.60 |
| Gill | Priest | Limnetic | 10 | 9 | 19 | 4.27 +/- 0.53 |
| Gill | Paxton | Benthic | 7 | 10 | 17 | 3.97 +/- 0.56 |
| Gill | Paxton | Limnetic | 10 | 10 | 20 | 4.40 +/- 0.37 |
| Gill | Little Quarry | Benthic | 11 | 9 | 20 | 4.01 +/- 0.38 |
| Gill | Little Quarry | Limnetic | 7 | 9 | 16 | 4.24 +/- 0.85 |
| **Gill total** |  |  | **55** | **57** | **112** | **4.13 +/- 0.58** |
| Liver | Priest | Benthic | 9 | 10 | 19 | 2.47 +/- 0.57 |
| Liver | Priest | Limnetic | 9 | 9 | 18 | 2.98 +/- 0.49 |
| Liver | Paxton | Benthic | 5 | 9 | 14 | 2.60 +/- 0.87 |
| Liver | Paxton | Limnetic | 11 | 10 | 21 | 2.86 +/- 0.35 |
| Liver | Little Quarry | Benthic | 9 | 7 | 16 | 2.49 +/- 0.91 |
| Liver | Little Quarry | Limnetic | 7 | 9 | 16 | 2.93 +/- 0.54 |
| **Liver** |  |  | **50** | **54** | **104** | **2.73 +/- 0.65** |

We extracted total RNA from gill and liver using the Zymo Quick-RNA Miniprep Plus Kit (Zymo, Irvine, California, United States), including the optional DNase treatment and eluting in 50 µl of nuclease-free water. RNA was quantified using the Qubit RNA BR Assay Kit (Invitrogen, Carlsbad, California, United States). We performed 3’-enriched mRNA sequencing library preparation using the MERCURIUS BRB-seq kit (Alithea Genomics, Épalinges, Switzerland). We used 230 and 210 ng of RNA as input for gill and liver, respectively. We used an epMotion 5070 automated pipetting system (Eppendorf, Hamburg, Germany) to transfer input RNA to the oligo(dT) plate and for pooling of first-strand synthesis products. All steps were conducted separately for each tissue (i.e. separate reaction plates, pools, sequencing lanes, etc.). To ensure evenness in sequencing depth among samples, we first performed shallow sequencing on an initial pool containing 3 µl from each reaction on a MiSeq Nano for gill and a iSeq 100 for liver (Illumina, San Diego, California, United States). Shallow sequencing was performed by the University of California San Diego Institute for Genomic Medicine’s Genomics Center in 28x90 bp format. Read depth per sample was calculated using the first several steps of our bioinformatic pipeline (see below). Using these depths, we adjusted the volume of cDNA from each reaction in each tissue’s final pool to normalize by number of reads. A small number of libraries were excluded from these pools due to poor sequencing performance. Each final pool was sequenced across two AVITI flowcells (Element Biosciences, San Diego, California, United States) using the 150-cycle kit (again in a 28x90 bp format). Deep sequencing was performed by the University of California Davis DNA Technologies Core.

### Read processing and differential-expression analysis

BRBseq libraries incorporate internal barcodes for sample multiplexing rather than conventional Illumina-style indices. We thus used the solo mode of STAR v2.7.11b (Dobin et al. 2013) to obtain sample-specific alignments to the *G. aculeatus* v5 reference genome (Nath et al. 2021), using the NCBI RefSeq annotations. STARsolo parameters were selected according to the Alithea BRBseq documentation. Gene-level counts were obtained using FastReadCounter v1.1 (Deplancke Lab 2023). For shallow sequencing, genome-wide counts per sample were obtained by modifying the attributes of the RefSeq gtf to consist of a single feature. The deep sequencing FastReadCounter output matrix was used as input to downstream differential expression analysis.

Prior to differential expression analysis, we constructed a PCA for each tissue from counts per million-normalized expression data to identify potential outlier samples. We identified one liver sample as an extreme outlier and excluded it from further analyses. We then used the R package DESeq2 v1.34.0 to perform differential expression analysis (Love et al. 2014). Each tissue was analyzed separately using all individuals across lakes and ecotypes with the following formula: ∼ sex + lake + ecotype + lake:ecotype. Therefore, dispersions were estimated across the entire set of fish within each tissue. Sexes were determined from whole genome resequencing data generated for a separate project. Lake-specific ecotype effects were extracted using the DESeq2 results() function on the fitted dds object. FDR-adjusted p-values were calculated using the Benjamini-Hochberg method (Benjamini and Hochberg 1995) after independent filtering as implemented in DESeq2’s defaults. For downstream methods using log_2_-fold changes (LFCs), we used the lfcShrink() function (type=”ashr”) to minimize the contribution of low count and/or noisy genes. Benthic was used as the reference level; thus, positive LFCs indicate greater expression in limnetic fish and negative LFCs indicate greater expression in benthic fish.

### Quantifying parallel transcriptional divergence

To assess the extent of parallelism (or lack thereof) of differential expression between ecotypes across lakes, we considered a gene to be parallel between two lakes if 1) it was significant at an FDR ≤ 0.05 and 2) its LFC sign matched (e.g. same direction of expression in both lakes). We categorized genes as two- or three-way parallel contingent on being directionally parallel between any lake pair or across all three lakes, respectively. These categories are mutually exclusive unless otherwise noted (i.e. two-way parallel genes do not include those that are also three-way parallel). Because requiring significance in both lakes is sensitive to differences in statistical power, we also evaluated directional agreement among genes significant at FDR ≤ 0.05 in at least one member of each lake pair. Within this set, we calculated the proportions with concordant versus opposing signs of the ashr-shrunken LFC and visualized the paired lake-specific LFCs. This analysis therefore assessed whether directional concordance extended to expression responses that did not reach significance in both lakes.

We conducted permutation tests to evaluate whether the observed extent of directional parallelism exceeded that expected by chance. To generate null distributions of pairwise and three-way overlaps, we performed 1,000 permutations in which, within each lake, we randomly reassigned the direction of differential expression among genes identified as significantly differentially expressed (FDR ≤ 0.05), while holding gene identity fixed. We then aggregated overlaps across lakes using the same procedure applied to the observed data. Because some genes are more likely to be detected as significantly differentially expressed, due to higher base mean expression or greater variance, for example, conditioning on significance and permuting only the sign of expression change avoids exchanging highly detectable genes with less detectable ones. This approach yields a more permissive null model and therefore a conservative test of directional parallelism. Accordingly, this null tests excess directional concordance conditional on the observed set of significant genes, rather than excess overlap in the identities of significantly differentially expressed genes.

At the functional level, we used gene set enrichment analysis (GSEA) to test for coordinated expression divergence in KEGG pathways (Subramanian et al. 2005; Kanehisa and Goto). Stickleback genes were assigned to reference KEGG pathways through their KEGG Orthology annotations.

After filtering KEGG annotations to retain interpretable biological categories (Supplementary Methods), we analyzed gill and liver separately and performed an independent GSEA for each of the three lakes, for a total of six population-by-tissue analyses. For each analysis, all genes with a valid NCBI Gene ID and non-zero expression were ranked by their ashr-shrunken LFC. GSEA was run in clusterProfiler v4.14.0 (Yu et al. 2012), retaining pathways represented by 5 to 5,000 tested genes. Significance was defined as a Benjamini-Hochberg-adjusted p-value less than 0.05 within each population-by-tissue analysis.

To quantify parallel functional enrichment, for each tissue we compared the GSEA results across the three lakes. Positive normalized enrichment scores (NES) indicated enrichment among genes with greater expression in limnetic fish, whereas negative NES indicated enrichment among genes with greater expression in benthic fish. A pathway was considered directionally parallel between two lakes when it was significant in both and had NES values with the same sign; three-lake parallelism required significance and concordant NES direction in all three lakes. For visualization and interpretation, we summarized the magnitude of enrichment with the mean NES across all three lakes and its among-lake range, focusing on pathways with directionally concordant enrichment in at least two lakes.

To ask whether parallel pathway enrichment arose from similar gene-level responses, we compared the ashr-shrunken LFCs of member genes among lakes. Within each tissue and pathway, we retained genes with an absolute LFC greater than 0.1 in at least one lake, thus excluding genes with uniformly negligible effects while retaining genes that responded strongly in only a subset of lakes. We then calculated Spearman correlations between lake-specific LFCs. For pathways parallel in two lakes, we used the correlation for the concordantly enriched lake pair; for pathways parallel in all three lakes, we averaged the three pairwise correlations. Higher correlations indicate greater conservation of the relative ordering of member-gene responses among lakes, whereas lower correlations indicate that similar pathway-level enrichment arose from different gene-level patterns.

### Evolutionary constraint and expression repeatability

If evolutionary constraint limits the range of viable regulatory responses available during adaptation, it may also increase the repeatability of gene-expression divergence across independently evolved populations. We therefore tested whether proxies of pleiotropic and sequence-level constraint predict the consistency of gene-expression responses, first across the transcriptome and then among genes within pathways showing parallel functional divergence.

To quantify the repeatability of gene expression responses across lakes at the whole-transcriptome level, we ranked all expressed genes by their ashr-shrunken LFC within each lake and scaled ranks from 0 to 1 as (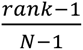, where N = total number of expressed genes) in each lake. For each gene, we calculated the absolute difference in scaled rank between each pair of lakes and averaged these differences across the three lake pairs to obtain scaled rank divergence *D_SR_*. Smaller *D_SR_* values indicate that a gene retained a more similar position within the transcriptome-wide distribution of expression responses across lakes. The expected absolute difference between two independently assigned scaled ranks approaches 1/3 as N becomes large. We therefore defined excess stability as:

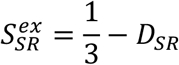

Positive excess stability indicates that a gene retained a more similar transcriptome-wide rank across lakes than expected, whereas negative values indicate greater rank divergence than expected.

Next, we adapted this metric to quantify the repeatability of member-gene responses within parallel pathways. This allowed us to ask whether proxies of evolutionary constraint predict repeated use of particular pathway constituents within pathways showing parallel functional divergence. The calculation of *D_SR_* was identical to the approach above, except that ranks were scaled relative to other genes within each pathway rather than across the full transcriptome. Because our definition of parallel pathways included both pathways showing parallel enrichment in only two lakes (e.g., Priest and Paxton but not Little Quarry) and pathways showing parallel enrichment across all three lakes, the calculation of *D_SR_* differed slightly between these cases. For two-way parallel pathways, *D_SR_* was simply the absolute difference in scaled rank between the two lakes showing parallel enrichment. For three-way parallel pathways, we calculated *D_SR_* as the mean absolute scaled-rank difference across all possible pairwise comparisons across lakes, as in the transcriptome-wide approach. Because pathways are finite and vary in size, the expected *D_SR_* under independent rank assignment depends on the number of genes in the pathway, N, and was calculated as:

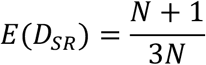

We therefore defined our pathway-level metric of repeatability, excess reuse, as

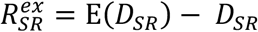

Positive excess reuse indicates that a gene retained a more similar rank relative to other members of the same pathway across lakes than expected under independent rank assignment, whereas negative values indicate greater pathway-relative rank divergence than expected.

### Proxies of evolutionary constraint

We assembled five proxies of evolutionary constraint, three closely linked to pleiotropy and two indicating constraint at the level of nucleotide sequence evolution. Our three pleiotropy metrics include expression breadth vs. specificity across tissues, KEGG pathway annotation degree, and gene co-expression network connectivity, which we explain below. The two metrics involving sequence evolution include the binary observation of nonsynonymous mutations and the dN/dS ratio between *G. aculeatus* and its congener *G. nipponicus*. As these five gene properties are not independent dimensions of constraint, their effects should therefore be interpreted as partially overlapping signatures of gene-level evolutionary constraint rather than as acting independently.

To characterize the degree of overlap among these gene properties, we calculated pairwise-complete Pearson correlations among the five focal constraint proxies and mean expression separately for gill and liver. Correlations used the transformed variables employed in the statistical models (see below), including log10-transformed KEGG annotation degree, dN/dS, and mean expression. Pairwise-complete observations allowed each comparison to retain all genes with data for that pair rather than restricting every comparison to the smaller set with valid dN/dS estimates.

#### Tissue specificity

Genes that are more broadly expressed across tissues are more likely to be pleiotropic because their products function in a wider range of cellular and physiological contexts (Watanabe et al. 2019). We estimated tissue specificity using the tau (*τ*) index originally described by Yanai et al. (2005). *τ* ranges from 0 to 1, with values near 0 indicating similar expression across many tissues and values near 1 indicating expression concentrated in a small number of tissues. We used public stickleback microarray data from NCBI’s Gene Expression Omnibus (GEO) series GSE34783, generated for Jones et al. (2012) which profiled seven adult tissues from marine and freshwater *G. aculeatus*: eye, gill, heart atrium and ventricle, hypothalamus, liver, pectoral muscle, and telencephalon.

Because the array was designed against an older stickleback assembly, we remapped array probes to the transcript set from the *G. aculeatus* reference used in this study. Probe sequences were aligned to transcript sequences with blastn-short (Camacho et al. 2009). We retained probes that mapped to a single gene and kept multi-mapping probes only when the top gene hit exceeded the second-best gene hit by at least 10 bitscore units. Probes without an unambiguous gene assignment under these criteria were excluded. For each retained probe, the processed GEO log2 expression ratio was averaged within each tissue and then converted back to a sample-to-reference ratio. For genes represented by multiple retained probes, we used the median expression ratio across probes within each tissue.

For each gene, *τ* was calculated across the seven tissues as:

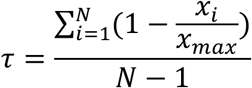

where *x_i_* is the gene’s expression estimate in tissue *i*, *x_max_* is the maximum expression estimate for that gene across tissues, and *N* is the number of tissues with non-missing expression estimates. For downstream models of excess reuse and excess stability, *τ* was centered and scaled prior to subsetting to genes included in each tissue’s transcriptome.

#### KEGG annotation degree

Genes annotated to a greater number of KEGG pathways may be more pleiotropic, as they are potentially involved in a greater number of cellular processes. While the annotation degree (number of KEGG pathways to which a gene is annotated) is also surely influenced by both the nested structure and variable granularity of available annotations and the uneven experimental effort across genes, it represents a simple and readily interpretable estimate of pleiotropy. For use in downstream statistics, we log_10_-transformed, centered, and scaled annotation degree.

#### Gene co-expression network connectivity

We quantified gene co-expression connectivity separately for gill and liver using weighted gene co-expression network analysis (WGCNA) (Langfelder and Horvath 2008). Genes were retained when they had at least 0.1 counts per million in at least 10% of samples, leaving 20,020 genes in gill and 17,447 genes in liver. Counts were variance-stabilized with DESeq2. To prevent connectivity from being driven by mean expression differences among sampling groups, we used the removeBatchEffect() function from limma v3.50.3 (Ritchie et al. 2015) to remove the additive effects of sex, lake, and ecotype from each tissue-specific expression matrix before network construction.

After evaluating soft-power thresholds across a range of values, we constructed signed WGCNA networks using soft-thresholding powers of 4 for gill and 10 for liver. For each gene, total connectivity (kTotal) was calculated using the intramodularConnectivity() function. We divided kTotal by the number of possible connections (the number of network genes minus one) and then centered and scaled the resulting values within each tissue. Higher standardized connectivity therefore indicates that a gene had stronger average positive co-expression with the rest of the tissue-specific network. Standardized connectivity was centered and scaled within each tissue prior to downstream statistical modeling.

#### Estimates of rates of sequence evolution

To quantify coding-sequence constraint, we identified single-copy orthologs between the *G. aculeatus* and *G. nipponicus* reference genomes (Yoshitake et al. 2022) and estimated gene-wide pairwise dN/dS using codeml from PAML v4.8 (Yang 2007). Detailed CDS filtering, orthology inference, codon-aware alignment, and codeml settings are provided in Supplementary Methods. After discarding genes with dN/dS estimates exceeding 10 or standard errors greater than 5, the final set contained 13,619 *G. aculeatus* genes.

We observed that when no non-synonymous mutations are observed, PAML reports a dN/dS value of 0.001. Rather than treat these data as continuous, we categorized genes as containing a non-synonymous mutation or not, allowing us to make use of all genes evaluated by PAML. We also retained the continuous dN/dS metric by subsetting to those genes with observed non-synonymous mutations, centering and scaling the values prior to statistical modeling.

### Statistical models and robustness analyses

We first tested associations between the five constraint proxies and transcriptome-wide excess stability. We used linear models fitted separately for gill and liver. Because the set of genes differed across predictors, each predictor was tested in a separate model. All models included log_10_ mean expression as a covariate; models involving dN/dS or the presence of a nonsynonymous substitution also controlled for alignment length. Continuous predictors were centered and scaled, and p-values for the five focal predictors were adjusted within each tissue using the Benjamini-Hochberg procedure.

We then tested the same associations using pathway-relative excess reuse, the within-pathway rank-stability metric defined above. Gill and liver were analyzed separately using linear mixed models fitted with lme4 and lmerTest (Bates et al. 2015; Kuznetsova et al. 2017). To isolate associations among genes within the same pathway, each focal predictor and model covariate was decomposed into a within-pathway deviation, defined as the gene’s value minus the pathway mean, and a pathway-mean component, calculated using the complete-case dataset for each model. Both components were included as fixed effects, and the within-pathway coefficient was treated as the focal test. Random intercepts for gene and pathway accounted for the occurrence of individual genes in multiple pathways and the non-independence of genes belonging to the same pathway. Predictor-specific covariates and multiple-testing correction followed the transcriptome-wide models, and models were fitted by restricted maximum likelihood.

We conducted two robustness analyses to determine whether these associations were artifacts of stability stemming from unresponsive genes (i.e. those with low LFCs) and/or associations between predictors and the lake-specific magnitudes of expression divergence. First, we repeated the whole-transcriptome models after retaining genes whose median absolute LFC across the three lakes was at least 0.322, equivalent to an approximately 1.25-fold expression difference in at least two lakes. Second, we fitted lake-specific models using the absolute shrunken LFC as the response and the same predictors and covariates used in the stability models.

Finally, as a complementary sensitivity analysis and an alternative to the continuous rank-based approach described above, we tested whether the five constraint proxies predicted directional concordance of benthic-limnetic expression divergence across all three lakes. Genes were classified as concordant when their ashr-shrunken LFC had the same non-zero sign in Priest, Paxton, and Little Quarry, and as discordant otherwise. We fitted predictor-specific logistic regression models separately for gill and liver using the same covariates as the transcriptome-wide excess-stability models. We repeated these models after retaining genes with an absolute LFC of at least log2(1.25) in one or more lakes. This excluded genes showing negligible expression shifts in all three lakes without conditioning inclusion on differential-expression significance. P-values for the five focal predictors were adjusted within each tissue and analysis using the Benjamini-Hochberg procedure, and effect sizes were expressed as odds ratios with 95% confidence intervals.

## Results

### Global patterns of gene expression variation

After excluding low-quality libraries and one extreme sample outlier, we estimated gene expression for 112 gill and 104 liver samples (Table 1). For gill, the mean number of reads per sample mapping to the *G. aculeatus* reference genome was 4.13M (SD = 0.58M, range = 1.99-5.39M). For liver, the mean number per sample was 2.73M (SD = 0.65M, range = 0.80-4.06M).

In both tissues, principal components analysis revealed strong divergence across the whole transcriptome with axes associated with sex and ecotype (Figure 1C-D). In contrast, variance was not partitioned by lake, and samples from different lakes were largely overlapping. In gill, strong separation across ecotype was observed along PC1, followed by sex along PC2. In liver, the first axis was strongly associated with sex, whereas the second was associated with ecotype. Curiously, five female benthics from Little Quarry clustered strongly with male benthics in the liver PCA (Figure 1D), though the reason for this clustering is unclear.

### Gene-expression divergence is strongly parallel but tissue dependent

For gill, tests for differential expression between limnetic and benthic ecotypes in Priest, Paxton, and Little Quarry revealed 6726 (36.0% of genes passing DESeq2 independent filtering), 2402 (13.9%), and 4654 (26.9%) significant genes (FDR <= 0.05), respectively (Figure 2A). The corresponding liver comparisons identified 2103 (14.4%), 1488 (10.5%), and 1979 (13.1%) significant genes in each lake.

**Figure 2.**
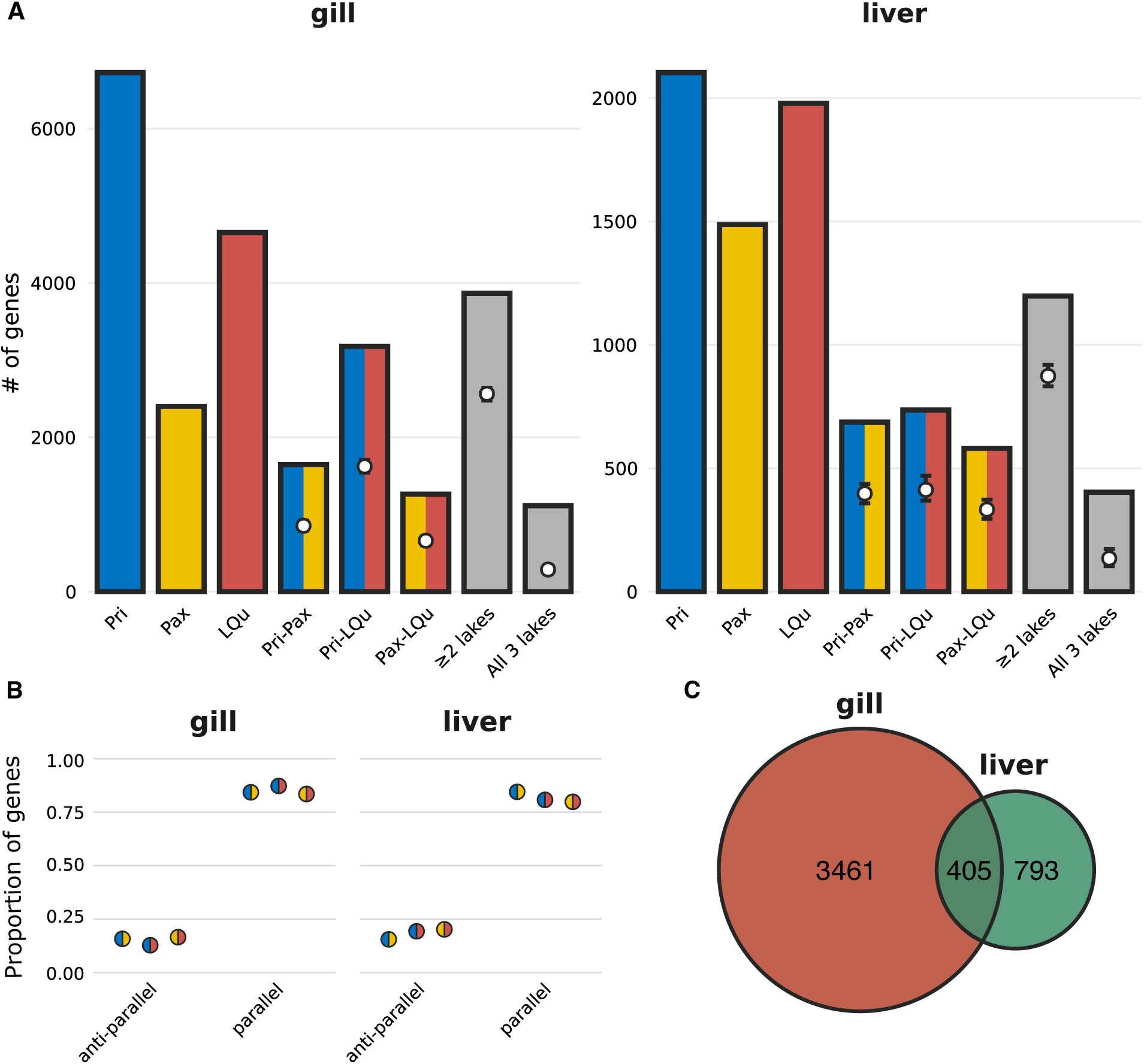
Evidence for widespread transcriptional parallelism among lakes. A) Results from differential expression analyses comparing ecotypes within each lake. First three columns represent the number of significantly differentially expressed genes in Priest (Pri), Paxton (Pax), and Little Quarry (LQu). Columns 4-6 represent pairwise overlaps of significant and directionally-concordant genes between all lake pairs. The final two columns represent the number of overlapping significant and directionally-concordant genes in at least one lake pair and among all three lakes. White points show the mean overlap expected under a permutation test in which LFC signs were randomly reassigned among significant genes within each lake, independently across lakes, while holding the identities and numbers of significant genes fixed. Error bars show the full range across 1,000 permutations. B) Proportions of genes exhibiting parallel and antiparallel expression divergence for each pair of lakes. For each pair, genes were included if they were significantly differentially expressed (FDR ≤ 0.05) in at least one lake. Genes were classified as parallel when LFC signs agreed between lakes and antiparallel when signs differed. Split-color points identify the two lakes in each comparison, using colors from Figure 1B. C) Overlap between genes showing repeated, directionally concordant expression divergence in gill and liver. Genes were included in a tissue-specific set when they were significantly differentially expressed in the same direction in at least one lake pair. The intersection contains genes meeting this criterion in both tissues; concordance of direction between gill and liver was not required.

Directionally parallel differential expression was pervasive. In gill, 3,866 genes (23.3%) exhibited significant, directionally concordant differential expression in at least two lakes, including 1,114 (6.4%) that were concordant in all three. In liver, the corresponding values were 1,198 (8.2%) and 403 (2.9%). Conditional on the observed identities of significantly differentially expressed genes, the number of shared genes with concordant LFC directions was 1.91-1.96-fold greater than expected under the sign-permutation null in gill and 1.73-1.78-fold greater in liver. Three-lake directional concordance was 3.87-fold and 2.99-fold greater than expected, respectively. Two-way parallelism was stronger for the Priest-Little Quarry pair than for the two pairs involving Paxton. This difference was especially pronounced in gill, stemming from the relatively small number of significantly differentially expressed genes in Paxton. All observed counts of directionally concordant shared-DE genes exceeded those obtained across 1,000 sign permutations (one-tailed empirical p = 0.001).

Directional agreement extended well beyond genes reaching significance in both lakes. Among genes significantly differentially expressed in at least one member of each lake pair, LFC directions were concordant for 83.5–87.2% of comparisons in gill and 79.8– 84.5% in liver (Figure 2B). The pairwise LFC distributions further showed that many genes significant in only one lake nevertheless had a same-direction, subthreshold response in the other lake (Figure S1). This indicates that parallel expression divergence extends beyond genes meeting significance thresholds in both lakes, with many subthreshold responses nevertheless changing in the same direction across independently evolved populations.

Of the genes showing significant, directionally concordant differential expression in at least two lakes, 3,461 were unique to gill, 793 were unique to liver, and 405 were shared between tissues (Figure 2C). However, while the majority (274, 67.7%) of these genes were differentially expressed in the same direction in gill and liver, 131 (32.3%) were antiparallel across tissues. Among the directionally concordant genes shared among two or more lakes, several benthic-biased genes had clear immune functions, including cd247, nkl.4, and ncf2, which encode a component of the T-cell receptor signaling complex, a cytotoxic effector, and a component of the phagocyte oxidative burst, respectively. Several genes overexpressed in limnetic gill but under-expressed in limnetic liver were associated with tissue remodeling and developmental signaling, including fbn1, adamts10, and fgfr2.

### Parallel functional divergence varies in underlying gene-level concordance

GSEA identified 97, 1, and 61 significantly enriched KEGG pathways in Priest, Paxton, and Little Quarry gill, respectively, and 65, 36, and 39 in liver. In gill, 45 pathways were significant and directionally concordant in two or more lakes (Figure 3A). Of these, 44 were parallel in two lakes, whereas only ribosome was significant in the same direction in all three. Because ribosome was the only significant pathway in Paxton gill, all 44 two-lake parallel pathways reflected the Priest-Little Quarry comparison. Of the 45 parallel gill pathways, 30 had positive NES values and were dominated by endocrine, neuronal, and cell-signaling functions. The 15 pathways with negative NES values were concentrated in core cellular functions such as DNA replication, oxidative phosphorylation, and ribosome.

**Figure 3.**
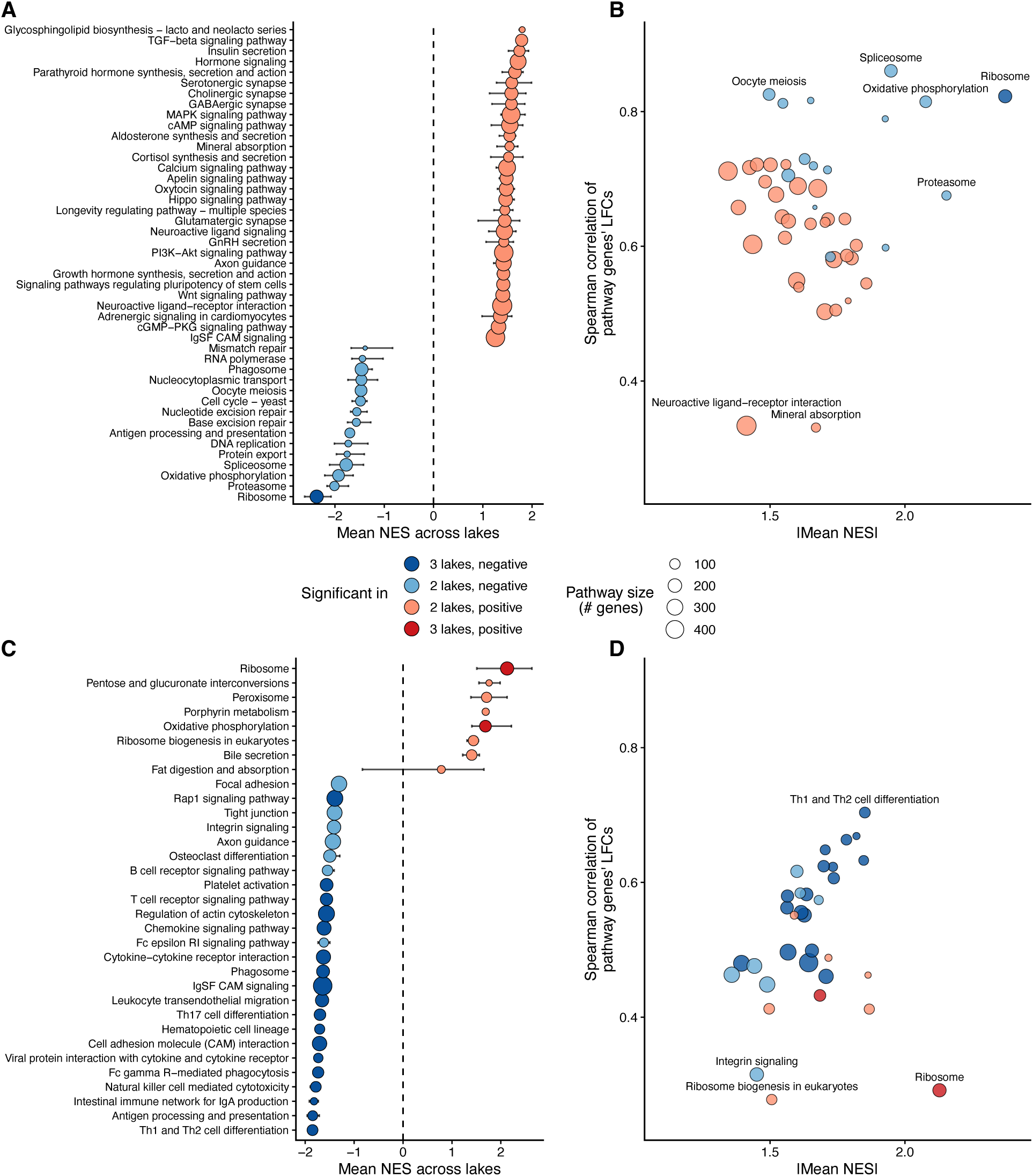
Extensive parallelism at the functional levels is underlain by variable gene-level parallelism. A,C) KEGG pathways exhibiting significant, directionally concordant enrichment in at least one lake pair in gill (A) and liver (C). The x-axis represents the mean normalized enrichment score (NES) across all three lakes. Point color indicates enrichment direction relative to benthic fish: positive NES indicates limnetic-biased enrichment, whereas negative NES indicates benthic-biased enrichment. Color intensity distinguishes pathways with concordant enrichment in two versus all three lakes. Point size represents pathway size, and horizontal error bars show the range of NES values across the three lakes. B,D) Relationship between enrichment magnitude and the repeatability of member-gene expression responses for KEGG pathways in gill (B) and liver (D). Points represent pathways showing significant, directionally concordant enrichment in at least two lakes. For pathways concordantly enriched in two lakes, the x-axis gives the absolute mean NES of the concordant lake pair and the y-axis gives the Spearman correlation between member-gene LFCs in that pair. For pathways concordantly enriched across all three lakes, the x-axis gives the absolute mean NES across the three lakes and the y-axis gives the mean of the three pairwise Spearman correlations. Correlations were calculated from ashr-shrunken LFCs for pathway-member genes with an absolute LFC greater than 0.1 in at least one lake. Point size represents pathway size, and point color indicates the number of concordant lakes and enrichment direction.

Although fewer pathways were parallel in liver than gill, three-lake parallelism was much more common: 20 of the 33 parallel pathways were significant with concordant direction in all three lakes, whereas 13 were parallel in two lakes (Figure 3C). Of the 33 parallel liver pathways, 25 had negative NES values, indicating greater expression in benthic livers, with especially consistent enrichment of immune, cell-adhesion, and cell-signaling pathways. Of these 25, 18 were significantly benthic-enriched in all three lakes, most with apparently immune-related functions. The eight parallel liver pathways with positive NES values were primarily associated with metabolism and digestive function. Ribosome and oxidative phosphorylation were significantly limnetic-enriched in all three lakes, opposite to what was observed in gill.

The direction of functional enrichment was strongly tissue dependent. As noted, ribosome was strongly and consistently enriched toward benthic expression in gill but toward limnetic expression in liver. Oxidative phosphorylation showed the same cross-tissue reversal. These contrasts show that repeated functional divergence between ecotypes did not represent a single organism-wide expression program, but instead involved distinct and sometimes opposing responses in gill and liver.

Parallel functional divergence was not always accompanied by strong correlations in the rank order of LFCs of constituent genes (Figure 3B,D), indicating that parallel pathway enrichment could occur despite relatively weak conservation of member-gene response ranks. Among gill pathways, several benthic-enriched pathways involved in core cellular processes such as oxidative phosphorylation and ribosome showed particularly strong conservation of member-gene responses. Further, correlations varied substantially even among pathways with similar enrichment magnitudes. Neuroactive ligand-receptor interaction and mineral absorption, for example, were repeatedly enriched toward limnetic expression but had correlations of only approximately 0.35. Thus, some parallel gill pathways were produced by strongly conserved relative responses among their member genes, whereas others reached similar pathway-level enrichment through less concordant gene-level changes.

Member-gene correlations were generally lower and more variable in liver (Figure 3D). The highest correlation was observed for Th1 and Th2 cell differentiation, and many of the three-lake benthic-enriched immune pathways had moderate-to-high correlations. This indicates that the repeated enrichment of immune functions was frequently accompanied by conservation of the relative ordering of gene responses across lakes. Other pathways showed much less gene-level concordance. Integrin signaling had one of the lowest correlations among benthic-enriched pathways, while ribosome biogenesis and ribosome also had correlations near 0.30. Notably, ribosome was one of the strongest and most consistently limnetic-enriched liver pathways, yet its member-gene correlation was among the lowest observed. This is opposite to what was observed in gill. Thus, not only is ribosome strongly and consistently enriched in opposite directions across tissues, but the extent of its underlying gene-level concordance is divergent as well, with gill exhibiting much stronger conservation and liver more flexibility. Taken as a whole, parallel functional divergence appears to arise through a mixture of conserved gene-level regulation and more flexible, pathway-level convergence involving different configurations of member-gene expression.

### Evolutionary constraint predicts cross-lake expression repeatability

The contrast between more conserved gene-level regulation and apparently flexible pathway-level convergence led us to ask how proxies of evolutionary constraint may predict the repeatability of gene expression responses. We first evaluated transcriptome-wide excess stability, then asked whether similar relationships were evident when repeatability was evaluated relative to genes within pathways showing parallel, directionally concordant enrichment.

The focal gene properties were not strongly correlated with one another (Figure S2). All pairwise correlations among the five constraint proxies were weak (|r| ≤ 0.174). The strongest correlation involving the mean-expression covariate was with network connectivity in both gill (r = 0.363) and liver (r = 0.478). Thus, broadly expressed, highly connected, and slowly evolving genes did not simply represent a single strongly correlated axis of gene properties.

At the whole-transcriptome level, several proxies of evolutionary constraint were significantly associated with excess stability (repeatability) in gill (Figure 4A). Excess stability decreased with increasing tissue specificity (β = −0.0093, p_adj_ < 10^-15^) and with the presence of an observed nonsynonymous substitution (β = −0.0228, p_adj_ < 10^-10^), while increasing with network connectivity (β = 0.0179, p_adj_ < 10^-15^). Continuous dN/dS was also negatively associated with excess stability (β = −0.0043, p_adj_ = 0.0016), whereas KEGG annotation degree remained unrelated to stability (β = 0.0003, p_adj_ = 0.855). Thus, broadly expressed, highly connected, and more slowly evolving genes generally occupied more similar positions in the transcriptome-wide distribution of expression divergence across lakes.

**Figure 4.**
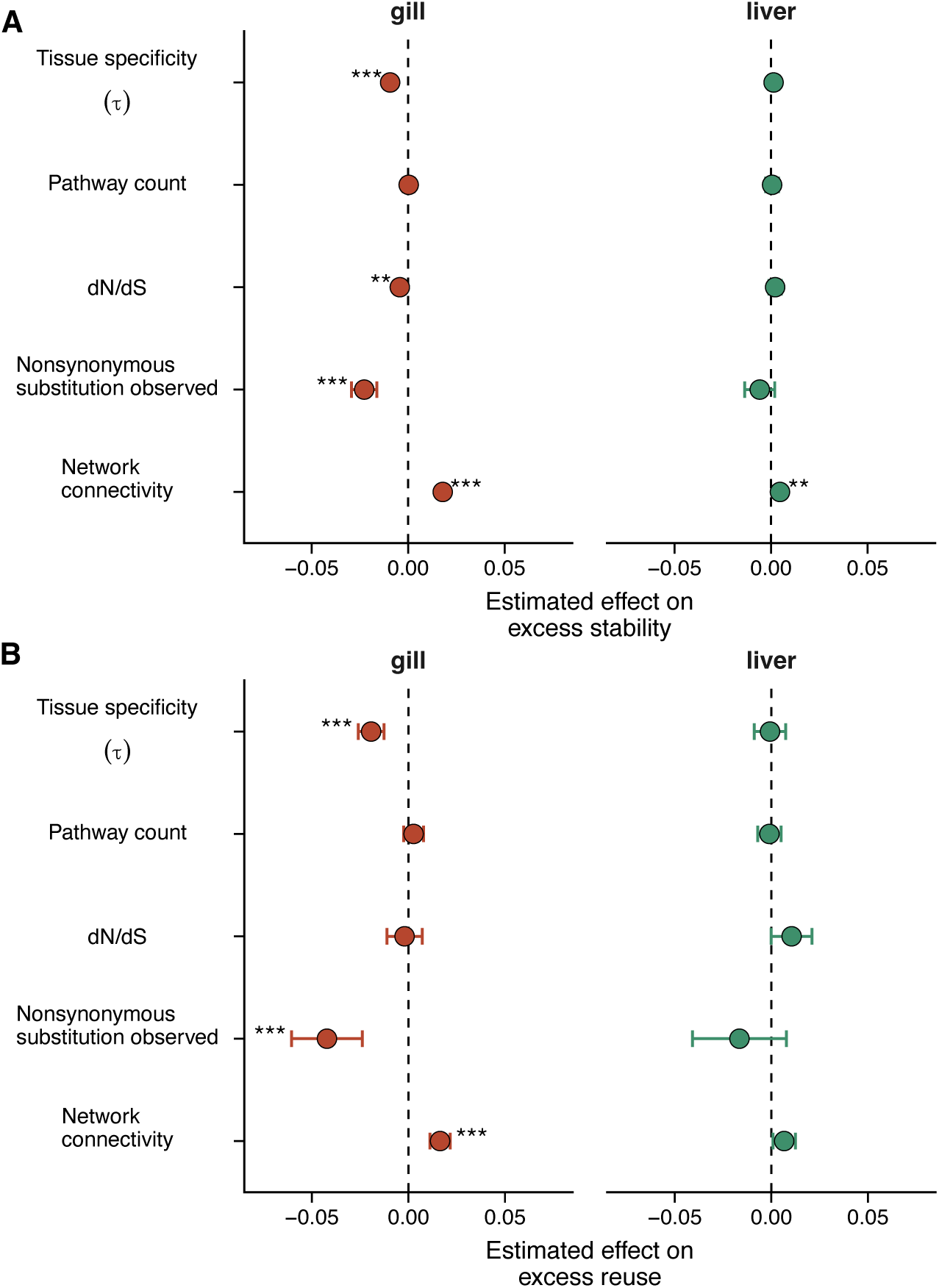
Proxies of evolutionary constraint predict the repeatability of expression divergence, particularly in gill. A) Estimated effects of five gene properties on transcriptome-wide excess stability. B) Estimated within-pathway effects of the same properties on excess reuse among genes belonging to KEGG pathways with parallel enrichment. Each predictor was tested in a separate model for gill and liver. Points represent estimated model coefficients, horizontal error bars represent 95% confidence intervals, and the dashed vertical line indicates no association. Positive coefficients indicate greater cross-lake expression repeatability. Continuous predictors were centered and scaled; the nonsynonymous-substitution coefficient contrasts genes with and without an observed nonsynonymous substitution. All models controlled for mean expression, and models involving coding-sequence evolution additionally controlled for alignment length. Continuous dN/dS effects were estimated only among genes with an observed nonsynonymous substitution. Pathway-level models included within-pathway and pathway-mean components of each predictor and covariate, as well as random intercepts for gene and pathway; the plotted coefficients represent the within-pathway effects. Asterisks denote Benjamini–Hochberg-adjusted significance within each tissue: * = p_adj_ < 0.05, ** = p_adj_ < 0.01, *** = p_adj_ < 0.001. Overall, broader expression, stronger coding-sequence constraint, and greater network connectivity predicted more repeatable expression responses in gill, whereas associations were substantially weaker in liver.

The corresponding transcriptome-wide relationships were weaker in liver (Figure 4A). Network connectivity was the only predictor significantly associated with excess stability (β = 0.00461, p_adj_ < 0.002); tissue specificity, KEGG annotation degree, dN/dS, and the nonsynonymous-substitution indicator were not significant (all p_adj_ ≥ 0.328). Thus, although highly connected genes exhibited modestly more stable expression divergence in liver, the broader constraint-associated pattern observed in gill was absent.

When associations of constraint proxies with repeatability were evaluated among genes within parallel pathways, three of the five predictors were significantly associated with pathway-relative excess reuse in gill after correction for multiple testing (Figure 4B). Tissue specificity had a negative effect (β = −0.0193, 95% CI = −0.0260 to −0.0126, p_adj_ = 4.02 x 10^-8^), indicating that broadly expressed genes retained more similar pathway-relative LFC ranks across lakes. Genes with an observed nonsynonymous substitution also had lower excess reuse (β = −0.0422, 95% CI = −0.0605 to −0.0239, p_adj_ = 1.09 x 10^-5^), whereas gene co-expression network connectivity was positively associated with excess reuse (β = 0.0164, 95% CI = 0.0112 to 0.0216, p_adj_ = 4.17 x 10^-9^). Neither KEGG pathway annotation degree (β = 0.0027, p_adj_ = 0.388) nor continuous dN/dS among genes with an observed nonsynonymous substitution (β = −0.0020, p_adj_ = 0.667) predicted pathway-relative reuse.

No predictor was significantly associated with pathway-relative excess reuse in liver after multiple-testing correction (all p_adj_ ≥ 0.126; Figure 4B). The estimated effects of co-expression network connectivity and the nonsynonymous-substitution indicator were in the same directions as in gill, but their confidence intervals overlapped zero. As with transcriptome-wide stability, within-pathway associations were substantially stronger in gill than in liver.

The transcriptome-wide associations were not driven solely by rank changes among genes with negligible expression divergence. After restricting the analysis to genes with a median absolute LFC of at least 0.322 across lakes, the negative effects of tissue specificity and an observed nonsynonymous substitution and the positive effect of connectivity remained significant in gill, while connectivity remained significant in liver (Figure S3). Continuous dN/dS in gill was similar in direction but fell just above the adjusted significance threshold (p_adj_ ≃ 0.052). Categorical logistic models of three-lake directional concordance generally recovered the patterns observed using the continuous rank-based approach, particularly the positive effect of connectivity in both tissues and the negative effect of an observed nonsynonymous substitution in gill, and these patterns were retained after excluding low-LFC genes (Figure S4). The main differences were positive effects of KEGG annotation degree in both tissues and positive effects of tissue specificity and dN/dS in liver, where their rank-based associations were weak and nonsignificant.

Lake-specific models of absolute LFC further showed that associations with repeatability were not generally explained by predictors simply producing larger expression differences (Figure S5). Tissue specificity in gill was positively associated with absolute LFC but negatively associated with stability, and the nonsynonymous-substitution indicator predicted stability without a corresponding association with absolute LFC. Connectivity was associated with both greater LFC magnitude and greater rank stability, so an effect mediated partly through response magnitude cannot be excluded for that predictor. In liver, all five predictors were positively associated with absolute LFC despite only connectivity predicting cross-lake excess stability. Together, these analyses indicate that the strongest constraint-associated pattern, particularly in gill, concerned the repeatability of genes’ relative expression responses across independent benthic-limnetic comparisons rather than simply the occurrence of larger expression differences.

## Discussion

While it is well-documented that repeated phenotypic evolution can progress through both parallel and idiosyncratic molecular trajectories, comparatively fewer studies have sought to explain this variation. Theoretical work and recent empirical examples highlight the potential importance of evolutionary constraint, including pleiotropy, in shaping molecular repeatability (Rennison and Peichel 2022; Battlay et al. 2024). Using repeated divergence of benthic and limnetic ecotypes of *G. aculeatus*, we find that genes under greater evolutionary constraint, as estimated by expression breadth across tissues, rates of nucleotide substitution, and coexpression network connectivity, exhibited more repeatable benthic-limnetic transcriptional divergence across lakes, particularly in gill. Importantly, the relationships between constraint and repeatability were not generally explained by constrained genes simply showing larger expression differences within lakes, suggesting a specific relationship between constraint and predictability rather than merely the magnitude of transcriptional divergence.

### Constraint predicts transcriptional repeatability

Classic evolutionary theory predicts that adaptation is shaped not only by the availability of beneficial mutations, but also by the range of phenotypic and molecular changes that can be tolerated. Pleiotropy has long provided a canonical example of such constraint. The Fisher-Orr model predicts a “cost of complexity,” whereby mutations affecting more phenotypic traits are less likely to contribute to adaptation because of their greater potential for deleterious pleiotropic effects (Fisher 1930; Orr 2000). More broadly, evolutionary constraint can arise from many features of gene function beyond pleiotropy and can be reflected over longer timescales in the strength of purifying selection acting on protein-coding sequence. Although pleiotropy can contribute to reduced rates of protein evolution, sequence-level constraint therefore represents a broader historical signature of restricted evolutionary change.

More recent work on repeated adaptation has highlighted a complementary consequence of constraint: by restricting the diversity of accessible evolutionary substrate, constraint may increase the repeatability of molecular evolution. Yeaman et al. (2018) explored this idea theoretically, arguing that repeated use of the same genes across independent lineages can reflect a restricted set of viable genetic routes arising from features of genetic architecture such as pleiotropy, epistasis, and limited redundancy in the genotype-phenotype map. Empirical support for a role of pleiotropy in shaping repeated gene use was provided by Rennison and Peichel (2022), who combined population genomic data from repeated stream-lake and benthic-limnetic ecotypic divergence events in stickleback with estimates of pleiotropy from prior studies. They demonstrated that genomic windows consistently identified as F_ST_ outliers across ecotype comparisons contained loci associated with intermediate levels of pleiotropy, as indicated by enriched overlap of QTL affecting multiple ecologically relevant traits and containing genes with higher co-expression network connectivity. Additional support for this view has since been provided by studies examining the propensity of genes to be reused during convergent evolution of repeated phenotypes across broader phylogenetic scales (Whiting et al. 2024; Barua et al. 2026).

These studies have primarily considered repeatability at the level of genomic loci, whereas gene expression provides an intermediate level of biological organization at which distinct genetic mechanisms may converge on similar molecular outcomes (Bolnick et al. 2018). Whether evolutionary constraint similarly predicts the repeatability of gene-expression divergence remains less clear, with previous studies yielding conflicting or null results (e.g. McGirr and Martin 2018; Durkin et al. 2024; Lai et al. 2025). Our results are most consistent with those of Lai et al. (2025), who likewise found that genes with greater expression breadth across tissues and higher co-expression network connectivity exhibited more predictable transcriptional responses during repeated adaptation. A complementary mechanism was identified by Koch et al. (2025) who used repeated adaptation during experimental evolution in *Tribolium* and found that highly connected genes experienced stronger indirect selection through genetic covariance with other expression traits, demonstrating how elevated coexpression network connectivity can facilitate rather than restrict adaptive transcriptional change. Our finding that transcriptional repeatability is also associated with stronger purifying selection at the coding-sequence level extends this pattern beyond pleiotropy-related proxies, linking repeated expression divergence to a deeper historical signature of evolutionary constraint.

Two features of our results are important for interpreting this relationship. First, support for an association between constraint and repeatability was substantially stronger in gill than in liver. Four of five proxies were associated with transcriptome-wide stability in gill and three were associated with pathway-relative reuse, whereas in liver only coexpression connectivity was significant at the transcriptome-wide level and no predictor remained significant for pathway-relative reuse after correction. However, several liver effect estimates were in the same direction as those observed in gill, suggesting that the weaker liver results may reflect a reduced or less readily detectable signal rather than a qualitatively different relationship. Second, these associations were not generally reducible to constrained genes having larger expression effects. In gill, tissue specificity was positively associated with absolute LFC but negatively associated with stability, while the presence of a nonsynonymous substitution predicted stability without a corresponding effect on absolute LFC; connectivity was associated with both larger effects and greater stability. In liver, all five proxies were associated with larger absolute LFCs despite only connectivity predicting cross-lake stability. Thus, the strongest constraint-associated signal, particularly in gill, concerns the predictability of relative expression responses rather than simply their magnitude. Because expression was measured in wild-caught fish, however, these associations do not by themselves establish that evolutionary constraint acted specifically on heritable regulatory divergence.

### Population-genetic context and molecular repeatability

Population-genetic processes can modify how features of genetic architecture translate into molecular repeatability. Battlay et al. (2024) explored this interaction using individual-based simulations of repeated adaptation under varying levels of migration, effect size, and pleiotropy. Consistent with classical expectations, deleterious pleiotropy could reduce genetic repeatability, particularly as mutations affected increasing numbers of traits. However, this effect was weakened in the presence of gene flow: migration-selection balance favored repeated contributions from loci capable of producing larger-effect alleles, even when those alleles incurred pleiotropic fitness costs. Thus, gene flow can counteract some of the constraining effects of pleiotropy by restricting adaptation to a smaller subset of loci with sufficiently large net beneficial effects.

Population size may provide another source of context dependence. Even closely related experimental studies in *Drosophila simulans* have recovered contrasting relationships between pleiotropy and transcriptional repeatability. Lai et al. (2025) found greater parallelism among more pleiotropic genes, whereas Ramirez-Lanzas et al. (2025) found that highly pleiotropic genes contributed disproportionately during the initial phase of adaptation but that more persistent and repeatable responses were biased toward genes with lower-to-intermediate pleiotropy. Ramirez-Lanzas et al. (2025) proposed the approximately 100-fold difference in population size between the experiments as one possible explanation, wherein stronger drift in smaller populations may restrict repeatable responses to variants with sufficiently large effects, but weak drift in very large populations permits selection on a more highly polygenic set of lower-effect changes. The small populations used in the aforementioned *Tribolium* experiment (120 breeding adults) are consistent with, but do not test, this possibility (Koch et al. 2025). Because population size was not experimentally manipulated in these studies, its role remains hypothetical.

These considerations are particularly relevant to benthic-limnetic stickleback, in which repeated ecological divergence has occurred in sympatry and is maintained despite incomplete reproductive isolation and ongoing gene flow. Hybridization and introgression continue between benthic and limnetic ecotypes in both Paxton and Priest lakes, with evidence for historically greater gene flow between ecotypes in Paxton (Gow et al. 2006; Schluter et al. 2025). Consistent with this history, Paxton exhibited the weakest benthic-limnetic transcriptional divergence in our dataset and was consequently less frequently represented in parallel expression responses shared across lakes, particularly in gill. Variation among lakes in the extent and history of gene flow may therefore contribute to variation in both the magnitude and repeatability of transcriptional divergence.

Demographic history should also shape repeatability through the genetic variation available to selection. Postglacial freshwater colonization was accompanied by founder effects and reduced genetic diversity relative to marine populations (Taylor and McPhail 2000), while repeated freshwater adaptation in stickleback is well known to draw heavily on standing genetic variation (Jones et al. 2012; Roberts Kingman et al. 2021). This combination of demographic filtering and shared variation may help explain the relatively high genetic repeatability of benthic-limnetic divergence: nearly half of QTL underlying parallel phenotypic differences were reused between the Paxton and Priest species pairs (Conte et al. 2015), and marine-benthic crosses similarly recovered overlapping QTL across independently derived benthic populations (Erickson et al. 2016). Preliminary analyses from our ongoing population-genomic work likewise suggest that benthic populations share substantial derived variation across lakes, raising the possibility that repeated selection has acted on a common pool of standing variation. Shared variation may therefore supply common evolutionary routes, while drift and gene flow determine which of those routes contribute to and persist during divergence.

Variation in these factors may also help explain why molecular repeatability differs among ecotypic contrasts within stickleback. Benthic-limnetic divergence shows relatively extensive QTL reuse, whereas repeated lake-stream divergence can be phenotypically parallel despite little reuse of the same QTL (Poore et al. 2023). Common-garden transcriptomes from independent lake-stream pairs likewise recovered only a small set of parallel expression differences (Hanson et al. 2017). Marine-freshwater transitions provide a contrasting axis in which recurrent selection on ancient standing variation is particularly well documented (Roberts Kingman et al. 2021), and parallel gill-expression divergence across four marine-freshwater pairs was enriched near adaptive genomic regions and for cis-regulatory control (Verta and Jones 2019). Comparisons across benthic-limnetic, lake-stream, and marine-freshwater divergence therefore provide an opportunity to test whether variation in population history, shared standing variation, and ongoing gene flow predicts molecular repeatability across both genetic and transcriptional levels. Comparable links between population history and molecular parallelism have also emerged in Arctic charr, lake whitefish, and cichlids (Derome et al. 2006; St-Cyr et al. 2008; Rougeux et al. 2017; Jacobs et al. 2020; Weber et al. 2021), emphasizing that the evolutionary routes available to selection are contingent on demographic history as well as properties of individual genes.

An additional feature of our results is that the extent of parallelism depended on the level of biological organization at which it was considered. Bolnick et al. (2018) emphasized that repeated evolution may vary across biological scales because many-to-one mapping allows distinct lower-level changes to generate similar higher-level outcomes. Evolutionary responses may thus become increasingly repeatable as they are integrated from individual variants and genes into pathways, physiological functions, and ultimately organismal phenotypes. Recent empirical work has begun to support this hierarchical model. In experimentally evolved *D. simulans*, (Lai et al. 2023) found that replicate populations with heterogeneous genomic and transcriptional responses nevertheless showed more parallel metabolomic responses, with different sets of responsive genes converging on similar biological functions. Cheek et al. (2026) explicitly compared parallelism across SNPs, genes, and pathways underlying repeated bill divergence in island scrub-jays, finding little overlap at the SNP or gene level but extensive parallelism at the pathway level. Our results reveal a similar pattern within the transcriptome itself. Parallel pathway enrichment could therefore coexist with substantial flexibility in member-gene response ranks, consistent with functional redundancy in the gene-level routes underlying similar pathway-level outcomes. Thus, functional redundancy can preserve flexibility in the molecular routes taken by independent populations even when selection repeatedly favors similar higher-level transcriptional outcomes.

### Parallel functional divergence reflects ecological specialization

Beyond explaining why some transcriptional responses are more repeatable than others, the identity of those repeated responses also provides insight into the physiological functions that repeatedly diverge between ecotypes. Despite extensive work on the genetic architecture and organismal phenotypes underlying benthic-limnetic divergence, its transcriptomic basis has received surprisingly little attention. To our knowledge, prior expression studies in these species pairs have been restricted largely to targeted measurements of cone opsins across lakes (Rennison et al. 2016) and a recent whole-brain RNA-seq study focused on female mate choice and sexual isolation (Keagy et al. 2024). Our data therefore provide the first comprehensive view of ecotype-associated transcription across two tissues and multiple benthic-limnetic pairs, revealing that many of the strongest functional differences are themselves repeated across lakes.

The most striking repeated functional differences involved core cellular and metabolic processes, but their direction was strongly tissue dependent (Figure 3). In gill, ribosome, oxidative phosphorylation, proteasome, DNA replication, and related cell-cycle pathways were repeatedly enriched toward benthic expression, whereas ribosome and oxidative phosphorylation were enriched in the opposite direction, toward limnetic, in liver, along with peroxisome, bile secretion, and lipid metabolism. This reversal suggests tissue-specific physiological specialization rather than a simple organism-wide increase in metabolic activity in either ecotype. The limnetic-biased hepatic signature is consistent with known ecological-performance differences. Blake et al. (2005) found that limnetics, which occupy the open-water habitat, exhibit greater prolonged swimming endurance than benthics. Sustained swimming in fishes relies heavily on aerobic metabolism and lipid fuels, with the liver playing a central role (Weber et al. 2016; van de Pol et al. 2017). Consistent with this interpretation, recent metabolomic work has also identified ecotype-associated divergence in the stickleback liver metabolome along the benthic-limnetic axis (Dinh et al. 2026), supporting a role for hepatic physiology in benthic-limnetic ecological differentiation.

Repeated divergence in liver also centered strongly on immune and cell-interaction functions, with benthics showing enrichment of pathways involved in lymphocyte signaling, phagosome, and leukocyte migration. Several benthic-biased genes, including cd247, nkl.4, and ncf2, likewise have direct immune functions. This pattern has clear ecological relevance because benthic and limnetic trophic niches expose fish to different parasite communities. Direct surveys of the Paxton and Priest species pairs found large, repeated differences in parasite burdens between benthics and limnetics (MacColl 2009), while across broader stickleback populations diet and trophic morphology predict parasite community composition and infection by trophically transmitted parasites (Stutz et al. 2014; Bolnick et al. 2020). Consistent with this ecological divergence, benthic and limnetic ecotypes also show parallel differences in MHC class II composition and allele number in Paxton and Priest lakes (Matthews et al. 2010). The repeated immune signatures observed here could therefore reflect an additional physiological dimension of ecological divergence, although they may represent both evolved differences and current differences in parasite exposure or immune state.

These functional interpretations should nevertheless be viewed at the tissue level. Because our data were generated from bulk gill and liver, some apparent expression differences may reflect shifts in cell-type composition rather than regulatory changes within the same cell types, particularly for immune-associated pathways. Thus, these data reveal highly repeatable tissue-level physiological differences between ecotypes, while leaving open the extent to which they reflect changes in cellular composition versus regulation within cell types. Because these fish were sampled from the wild, the observed expression differences likely integrate heritable regulatory divergence and plastic responses to ecotype-specific environments. Replication across independently evolved lake pairs makes purely idiosyncratic local environmental effects unlikely to explain the pervasive directional concordance, but it does not distinguish genetically encoded expression divergence from environmental responses that recur across ecologically similar benthic and limnetic niches. Thus, our data reveal highly repeatable ecotype-associated transcriptional divergence while leaving unresolved the relative contributions of evolved regulatory differences, plasticity, and shifts in cellular composition.

## Conclusion

Together, these results suggest that the predictability of transcriptional divergence depends on both the constraints acting on individual genes and the level of biological organization at which parallelism is measured. Parallel benthic-limnetic divergence was widespread, but the underlying gene-level responses ranged from conserved to flexible even when higher-level functional outcomes were similar. This combination points to an evolutionary landscape in which some transcriptional routes emerge repeatedly, while redundancy in regulatory and functional systems preserves multiple paths to similar physiological states. In this stickleback system, that balance may be further conditioned by population history, shared standing variation, and gene flow, helping explain why parallelism can be extensive without being uniform across genes, pathways, tissues, or lakes.

A key next step is to determine the evolutionary and regulatory mechanisms underlying these repeated expression phenotypes. Comparisons with contemporary marine populations can help polarize benthic-limnetic expression differences, revealing which ecotype has diverged further from the ancestral state and whether particular classes of genes are associated with more ancestral versus more derived stages of transcriptional divergence. Integrating genomic and transcriptomic variation will further make it possible to ask whether repeated expression differences arise through reuse of shared regulatory mechanisms or through distinct genetic routes. More broadly, extending this framework across different axes of adaptive divergence in stickleback should help reveal how evolutionary predictability emerges from the interaction between genetic architecture, demographic history, and the ecological context of adaptation.

## Supplementary Figures

**Figure S1.**
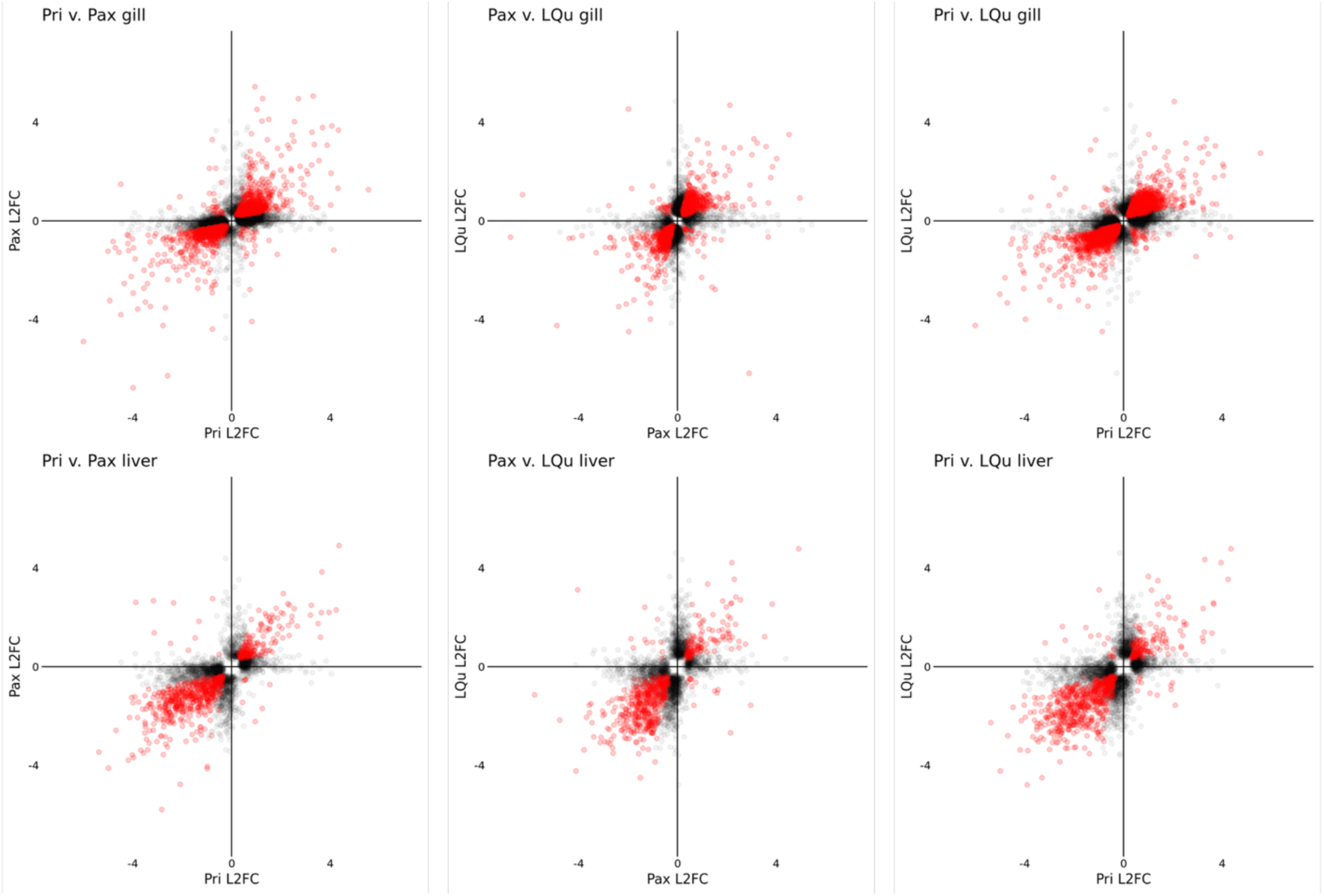
Pairwise expression divergence among lakes. Pairwise comparisons of ashr-shrunken LFCs for genes significantly differentially expressed (FDR ≤ 0.05) in at least one lake of each pair. Red points were significant in both lakes; black points were significant in only one.

**Figure S2.**
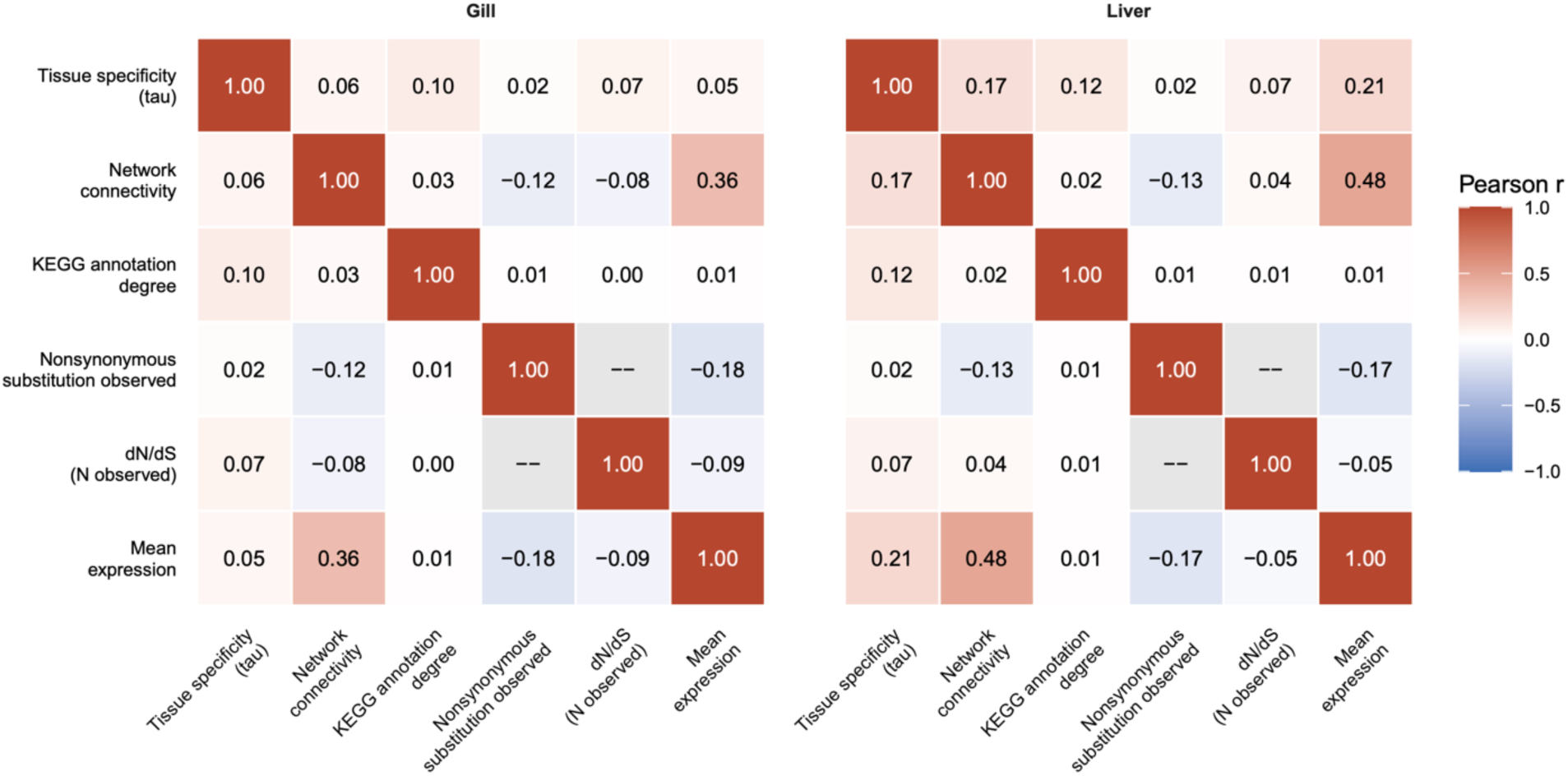
Correlations among focal gene properties. Pairwise-complete Pearson correlations among the five constraint proxies and mean expression, calculated separately for gill and liver.

**Figure S3.**
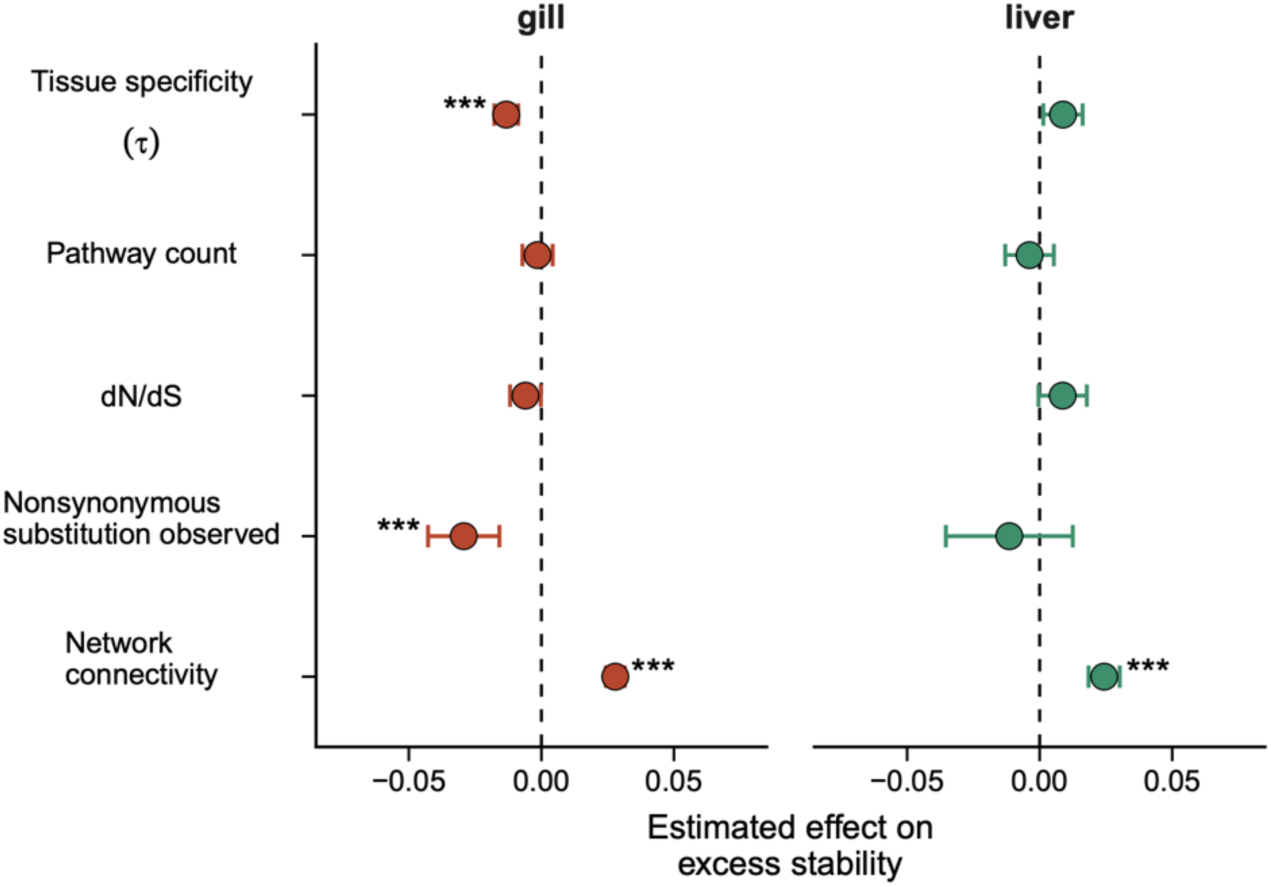
Excess-stability models excluding genes with negligible expression divergence. Estimated effects of the five gene properties on transcriptome-wide excess stability after retaining genes with a median absolute LFC ≥ 0.322 across lakes. Points show model coefficients and error bars show 95% confidence intervals. Asterisks denote BH-adjusted significance: * = p_adj_ < 0.05, ** = p_adj_ < 0.01, *** = p_adj_ < 0.001.

**Figure S4.**
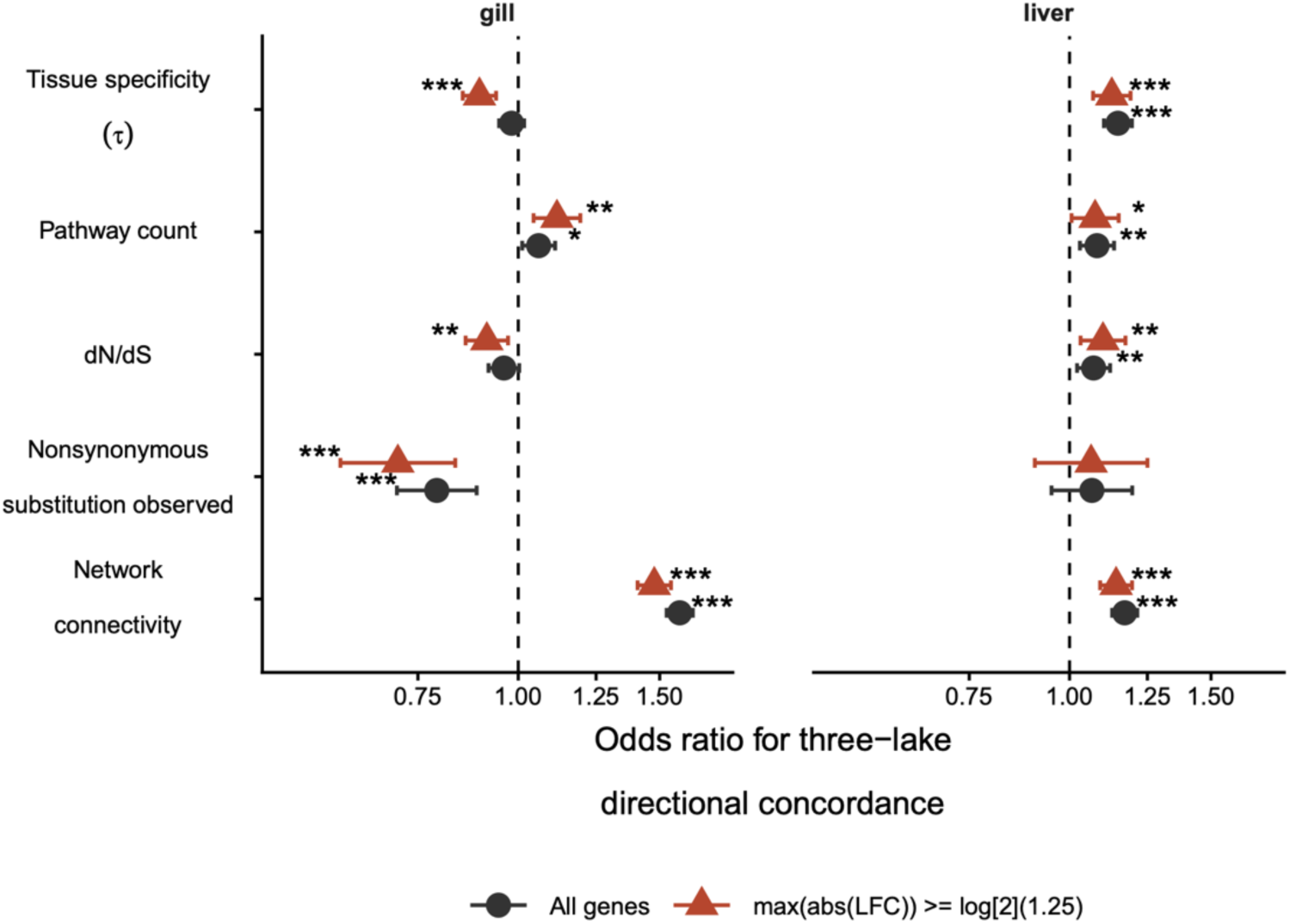
Categorical sensitivity analysis of directional concordance. Odds ratios from logistic models predicting whether genes had the same LFC sign in all three lakes. Results are shown for all genes and after requiring an absolute LFC ≥ log₂(1.25) in at least one lake. Error bars show 95% confidence intervals; asterisks denote BH-adjusted significance.

**Figure S5.**
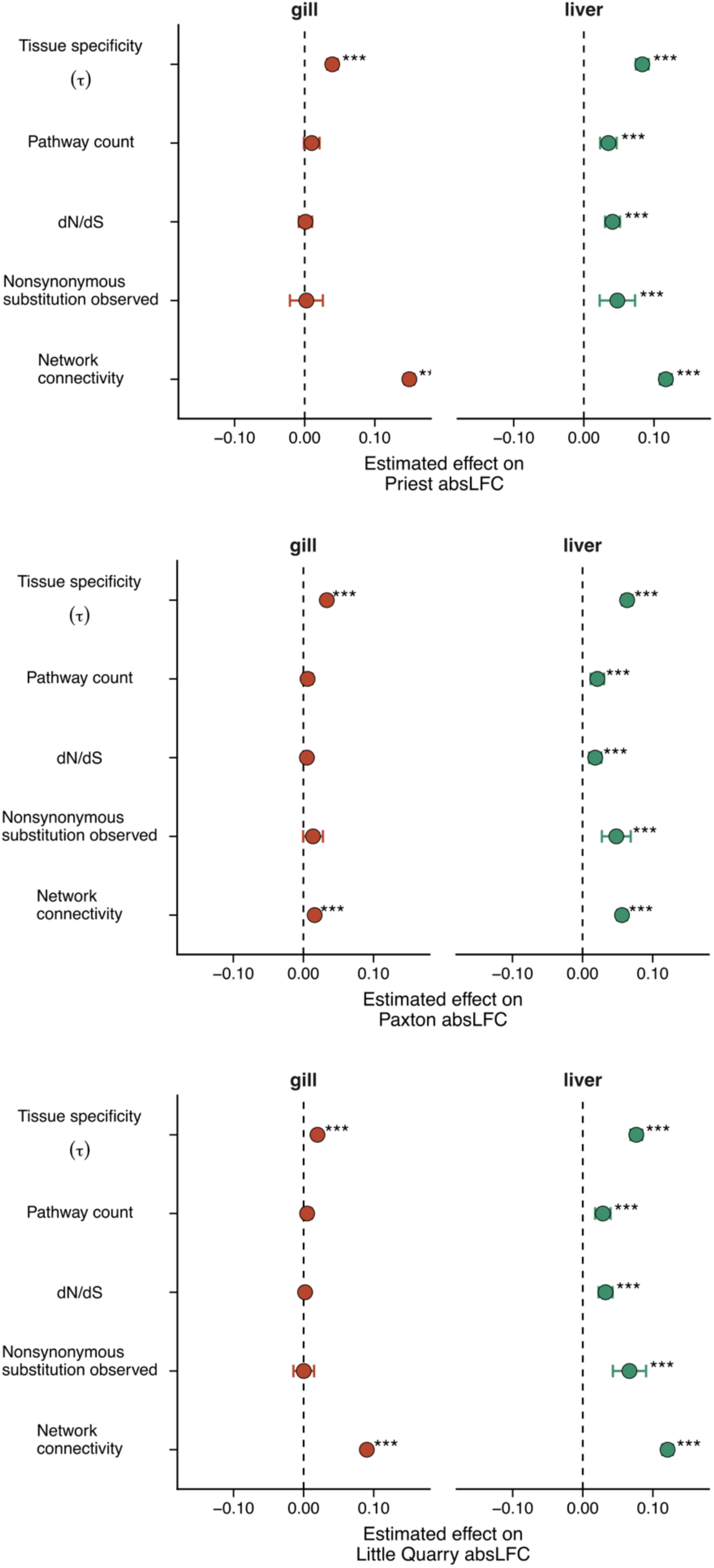
Associations between gene properties and expression-divergence magnitude. Estimated effects of the five gene properties on absolute LFC in Priest (A), Paxton (B), and Little Quarry (C). Points show model coefficients and error bars show 95% confidence intervals. Asterisks denote BH-adjusted significance.

## Supplementary Methods

The principal sampling, laboratory, and analytical procedures are described in the main Methods. Here, we provide additional implementation details for read processing, pathway analysis, construction of the constraint proxies and rank-based responses, and statistical sensitivity analyses.

### Read processing and differential-expression analysis

As described in the main Methods, reads were aligned to the G. aculeatus version 5 reference genome (GCF_016920845.1) with STAR v2.7.11b in STARsolo mode and counted with FastReadCounter v1.1. STARsolo was configured for a 14-bp sample barcode beginning at position 1 and a 14-bp UMI beginning at position 15 of the 28-bp barcode read. The MERCURIUS BRB-seq barcode list was supplied as the whitelist, libraries were treated as forward stranded, cell filtering was disabled, and only uniquely mapping reads were retained. UMI deduplication was not applied in STARsolo or during subsequent counting. For shallow sequencing, the RefSeq GTF was modified so that the annotated genome was represented as a single feature, providing one count per sample for pool balancing. Table 1 read-depth summaries were calculated from these counts after restricting the barcode columns to the final analytical sample set.

The preliminary outlier PCA used library-size-normalized counts after removal of genes with zero counts across the tissue dataset; genes were centered and scaled before PCA. The PCAs shown in Figure 1 were generated separately from the final DESeq2 objects using the variance-stabilizing transformation with blind=TRUE and all 28,430 rows in the count matrix.

Gill and liver were fitted separately in DESeq2 using the joint model described in the main Methods. Benthic and Little Quarry were the reference ecotype and lake, respectively. The Little Quarry limnetic-versus-benthic effect was therefore the ecotype main effect, whereas the Priest and Paxton effects were obtained by adding the appropriate lake-by-ecotype interaction to that main effect. Lake-specific results were extracted using alpha=0.05 and the default DESeq2 independent-filtering procedure. LFC shrinkage used type=“ashr” and the corresponding unshrunken lake-specific result object. The baseMean subsequently used as a covariate was the mean normalized count across the complete tissue-specific joint model and was therefore identical among the three contrasts derived from that model.

### Gene-level directional parallelism

Gene-level parallelism was classified as described in the main Methods. Pair-specific counts included genes that also met the three-lake criterion. In the aggregate categories, however, two-lake genes were defined as those parallel in at least one lake pair but not across all three lakes, making the two- and three-lake categories mutually exclusive.

For the sign-permutation test, positive and negative labels were shuffled independently within the sets of genes significant in Priest, Paxton, and Little Quarry. Thus, each permutation retained the identities and number of significant genes and the number of positive and negative effects in each lake. Pairwise, two-lake, and three-lake counts were then recalculated using the observed-data definitions. One-tailed empirical P values were calculated as [1 + number of null counts greater than or equal to the observed count]/1,001. The white points in Figure 2A show the means of the 1,000 null replicates, and their error bars show the complete minimum-to-maximum range. As emphasized in the main Methods, this tests directional concordance conditional on the observed significant-gene identities rather than excess overlap in the identities themselves.

The pairwise analysis in Figure 2B and Figure S1 instead used the union of genes significant in at least one member of each lake pair. Genes with a missing or exactly zero shrunken LFC in either lake were excluded because their direction was undefined. The tissue overlap in Figure 2C was constructed after independently identifying genes with repeated, same-direction significant divergence within gill and liver; agreement in the direction of divergence between tissues was not required for membership in the intersection.

### KEGG mapping, pathway filtering, and GSEA

RefSeq gene names were connected through a local cross-reference table to KEGG *G. aculeatus* identifiers and numeric NCBI Gene IDs. KEGG genes were then joined to KEGG Orthology identifiers and KO-linked reference pathways carrying the “map” prefix. NCBI Gene IDs were used for GSEA.

We excluded broad overview and chemical-transformation maps, categories labeled as human disease or drug development, and selected disease and drug-resistance categories that were not readily interpretable in stickleback. The manually excluded pathways were map01100, map01110, map01120, map01200, map01210, map01212, map01220, map01230, map01232, map01240, map01250, map01310, map01320, map01010, map01060-map01066, map01070, map04930, map04932-map04934, map04936, map04940, map04950, map01501-map01503, and map01521-map01524. We additionally removed all identifiers beginning with map05 or map07. After removal of incomplete mappings, 335 candidate pathways remained before applying the analysis-specific gene-set-size filter.

The six lake-by-tissue GSEA runs described in the main Methods used genes with a valid NCBI Gene ID and nonmissing nominal DESeq2 p-value, ranked from positive to negative by ashr-shrunken LFC. No significance or LFC threshold was applied before ranking. In clusterProfiler v4.14.0, TERM2GENE and TERM2NAME were supplied from the filtered mapping and pvalueCutoff=1, minGSSize=5, and maxGSSize=5,000 were used. Remaining settings followed the package defaults, including exponent=1, Benjamini-Hochberg adjustment, and the fgsea implementation.

For each tissue, lake-specific pathway results were joined before parallelism was classified. Pathways significant with the same NES sign in all three lakes were assigned three-lake parallelism. Those with at least one significant, same-direction pair but not the three-lake pattern were assigned two-lake parallelism. Consequently, the parallelism classification is not simply the number of lakes in which a pathway was significant; for example, a pathway significant in two lakes with opposing NES signs was not classified as two-lake parallel. For Figure 3A and C, the point estimate and interval were the mean and full range of all three NES values, including the NES from a lake in which the pathway was not significant.

The member-gene correlations in Figure 3B and D used duplicate-free pathway assignments and were not restricted to GSEA leading-edge genes. Within each tissue and pathway, the same union of genes with absolute LFC greater than 0.1 in at least one lake was used for all relevant pairwise correlations. Correlations were treated descriptively without a separate correlation P-value threshold.

### Construction of the evolutionary-constraint proxies

#### Tissue specificity

The seven-tissue tau index was calculated from the public microarray dataset described in the main Methods. The processed series contained 68 samples: ten each for eye, gill, heart, hypothalamus, and telencephalon and nine each for liver and pectoral muscle. Probe sequences were aligned to GCF_016920845.1 transcripts with blastn using task=blastn-short, an E-value threshold of 1e-8, dust filtering disabled, and a maximum of 50 target sequences per probe. For multi-gene probes, the best transcript hit for each candidate gene was first selected by bitscore, E-value, percent identity, and alignment length. A probe was retained only when the best gene exceeded the second-best gene by at least 10 bitscore units. Tissue means of the processed log2 sample-to-reference ratios were converted to ratio space, and multiple probes for a gene were summarized by their tissue-specific median. The focal tau value included all seven tissues.

#### KEGG annotation degree

Annotation degree was the number of distinct retained KEGG reference pathways assigned to a gene after collapsing duplicate gene-pathway pairs. Counts were log10 transformed and standardized. This measure was interpreted as annotation breadth rather than a direct count of biological functions.

#### Coexpression connectivity

Following the filtering and covariate removal described in the main Methods, counts were variance-stabilized with blind=FALSE. Sex and lake were supplied to limma’s removeBatchEffect as batch factors, and ecotype was supplied as the non-intercept column of an additional design matrix. Candidate powers of 1-10 and even powers from 12-40 were evaluated; powers 4 and 10 were selected for the gill and liver signed networks, respectively. WGCNA blockwiseModules was run with networkType=“signed”, TOMType=“signed”, minModuleSize=30, deepSplit=2, reassignThreshold=0, mergeCutHeight=0.25, maxBlockSize=40,000, numericLabels=TRUE, and pamRespectsDendrogram=FALSE. Signed adjacency was [(1 + correlation)/2]^power. The focal kTotal metric was the sum of a gene’s adjacency weights to all other network genes, scaled by the number of possible partners.

#### Coding-sequence evolution

Annotated coding sequences were extracted for G. aculeatus and *G. nipponicus* with gffread. We retained transcripts beginning with a canonical start codon, translated them with EMBOSS transeq, truncated peptides at the first stop codon, and selected the longest retained isoform per gene while preserving its nucleotide sequence. Single-copy orthologs were identified with OrthoFinder v2.4.1 using DIAMOND in ultra-sensitive mode, MAFFT, and FastTree. Each ortholog pair was realigned at the amino-acid level with MAFFT using --globalpair --maxiterate 1000, and PAL2NAL was used to create codon-aware PAML alignments.

Each pair was analyzed independently with codeml from PAML v4.8 using pairwise mode (runmode=-2), codon sequences (seqtype=1), the standard genetic code (icode=0), a single gene-wide omega (model=0 and NSsites=0), estimated omega with an initial value of 0.5 (fix_omega=0), FMutSel codon frequencies (CodonFreq=7 and estFreq=0), no among-site rate variation (fix_alpha=1 and alpha=0), and getSE=1. We retained estimates with dN/dS less than 10 and dN/dS standard error less than 5; no additional alignment-length or dS filter was applied. Alignment length was log10 transformed and standardized for use as a covariate. Values at the codeml floor of 0.001 were classified as having no observed nonsynonymous substitution. Continuous dN/dS models were restricted to genes above this floor and used log10-transformed, standardized dN/dS.

### Rank-based expression repeatability

The transcriptome-wide response described in the main Methods used genes with complete shrunken DESeq2 result rows in all three lake contrasts, including nonmissing LFCs and adjusted P values. This yielded 17,328 genes in gill and 14,150 in liver; no differential-expression significance filter was imposed. Average ranks were assigned to ties. The null expectation of two-thirds used for excess stability is the continuous large-N expectation; its difference from the exact finite-rank expectation is negligible for these transcriptome-wide sets.

For pathway-relative reuse, gene LFCs in each pathway and lake were first multiplied by the sign of that pathway’s NES in the lake. This placed the expression direction supporting pathway enrichment at the same end of the rank distribution across lakes before member genes were ranked. Pathway membership was defined from the retained KEGG map before merging any focal predictor, preventing predictor availability from defining the response. Forty-five gill pathways and 33 liver pathways met the parallel-enrichment criterion. Before predictor-specific missing-data filtering, the pathway tables contained 6,056 gene-by-pathway observations representing 2,941 unique genes in gill and 3,775 observations representing 1,832 genes in liver. Genes occurring in multiple pathways contributed one observation per pathway and could have different reuse values because their comparison sets differed.

For a pathway with n eligible genes, the expected absolute difference between two independently matched discrete ranks scaled from zero to one is (n + 1)/(3n), giving expected reuse (2n - 1)/(3n). Analytical excess reuse was observed reuse minus this pathway-size-specific expectation.

### Statistical models and sensitivity analyses

The predictor-specific models and covariates are described in the main Methods. Transcriptome-wide models were ordinary linear models, whereas pathway-relative models were linear mixed models fitted by restricted maximum likelihood. For the latter, within-pathway deviations and pathway means were calculated using the exact complete-case dataset for each model. The focal test was the within-pathway predictor coefficient. Models included crossed random intercepts for gene and pathway. Model-based 95% confidence intervals were reported, and the five focal p-values were adjusted by the Benjamini-Hochberg procedure separately within each tissue.

The gene-property correlation heatmaps used pairwise-complete Pearson correlations separately for gill and liver. Each cell therefore used all genes available for that pair rather than a common complete-case set. Continuous dN/dS was defined only among genes with an observed nonsynonymous substitution, making its correlation with the binary substitution indicator undefined.

For the sensitivity analysis restricted to genes with median absolute LFC of at least 0.322, transcriptome-wide ranks and excess-stability values were calculated before applying the threshold and were not recomputed within the reduced dataset. Transformations, covariates, and multiple-testing correction otherwise matched the primary models.

In the categorical sensitivity analysis, genes with three positive or three negative shrunken LFCs were classified as directionally concordant. Mixed signs or an exact zero in any lake were classified as discordant. The second analysis set required a maximum absolute LFC of at least log2(1.25) in one or more lakes; it did not select genes by differential-expression significance. Odds ratios and 95% Wald confidence intervals were obtained by exponentiating the focal log-odds coefficient and its coefficient plus or minus 1.96 standard errors. The five focal P values were adjusted separately within each tissue and analysis set. Lake-specific absolute-LFC models used the same predictor-specific covariates and restrictions as the primary stability models, with adjustment performed separately within each tissue and lake.

